# IS-Seq: a bioinformatics pipeline for integration sites analysis with comprehensive abundance quantification methods

**DOI:** 10.1101/2023.02.06.527381

**Authors:** Aimin Yan, Cristina Baricordi, Quoc Nguyen, Luigi Barbarossa, Mariana Loperfido, Luca Biasco

## Abstract

Integration site (IS) analysis is a fundamental analytical platform for evaluating the safety and efficacy of viral vector based preclinical and clinical Gene Therapy (GT). A handful of groups have developed standardized bioinformatics pipelines to process IS sequencing data, to generate reports, and/or to perform comparative studies across different GT trials. Keeping up with the technological advances in the field of IS analysis, different computational pipelines have been published over the past decade. These pipelines focus on identifying integration sites (IS) from single-read sequencing or paired-end sequencing data either using read-based or using sonication fragment-based methods, but there is a lack of a bioinformatics tool that automatically includes unique molecular identifiers (UMI) for IS abundance estimations and allows comparing multiple quantification methods in one integrated pipeline. Here we present IS-seq a bioinformatics pipeline that can process data from paired-end sequencing of both old restriction sites-based IS collection methods and new sonication-based IS retrieval systems while allowing the selection of different abundance estimation methods, including read-based, Fragment-based and UMI-based systems. We validated the performance of IS-Seq testing it against the most widely used analytical workflow available in the literature (INSPIIRED) and using different scenarios. Lastly, by performing a comprehensive wet-lab assessment of our IS-Seq pipeline we could show that in clinically relevant scenarios, UMI quantification provides better accuracy than the currently most widely used sonication fragment counts as a method for IS abundance estimation.

## Introduction

Integration site analysis (ISA) plays a key role for the monitoring of clonal dynamics of engineered cells in viral vector-based gene therapy. Recent developments in wet-lab protocols for insertion site retrieval and sequencing technologies have substantially expanded the reach and resolution of ISA. Concomitantly, the constant evolution of ISA techniques called for the development of bioinformatic tools for processing the sequencing results and automation of insertion sites mapping and quantification. Among the main computational pipelines developed over the past years, we can list IntegrationSeq/Map(Giordano et al., 2007), SeqMap(Peters et al., 2008) and SeqMap2.0(Hawkins et al., 2011), QuickMap(Appelt et al., 2009), MAVRIC(Huston et al., 2012), VISPA(Calabria et al., 2014), VISA(Hocum et al., 2015), GeIST(La Fave et al., 2015), the most widely used INSPIIRED(Berry et al., 2017; Sherman et al., 2017),VISPA2 (Spinozzi et al., 2017) and VSeq-Toolkit(Afzal et al., 2020). The general structure of these pipelines is based on processing and excluding a mosaic of oligos primer, adapters, sample barcodes, viral Long Terminal Repeat (LTR) sequences in order to release the host genomic DNA which will then be aligned to the host reference genome for the retrospective identification of the insertion sites. At the same time, specific bioinformatic tools embedded in these pipelines collect information relative to the abundance of each insertion site under the assumption that these will be reflective of the size of the relative clone contained in the original sample.

It is well known that the use of sequencing reads for abundance quantification can be, in certain scenarios, highly biased by the uneven PCR amplification of vector-genome junctions (Gillet et al., 2011). Deduplication can help removing PCR duplicates, but it has been shown that such process based exclusively on the use of mapping coordinates can also eliminate erroneously many usable reads (Fu et al., 2018). We have discussed previously why counting sonication fragments also has theoretical limitations in certain instances when it comes to accurate clonal abundance estimation (Biasco, 2017). To address this issue, Berry et al. developed a statistical inference approach (embedded in the sonicLength R package) based on deriving a likelihood function using the observed Fragment length-count data and applying a hybrid expectation–maximization algorithm to reach a Maximum Likelihood Estimation (MLE) of abundance estimation of vector-genome junctions (Berry et al., 2012). They showed that MLE based on Fragment data could indeed improve the quantification accuracy of ISA. In another work, Firouzi et al. showed that the use of a Unique Molecular Identifiers (UMI) tag system could further improve abundance estimation because of the high diversity of the UMI repertoire (Firouzi et al., 2014), although their analyses were limited to two clonal dilution points and their pipeline did not formally integrate the different quantification methods. Later, Rosewick et al developed an improved pipeline by combining Fragment length and UMI tag for abundance quantification based on a linear amplification mediated (LAM-)PCR method (Rosewick et al., 2020). Wells et al. combined experimental procedures with bioinformatics analysis to reduce the number of PCR/sequencing artifacts upon integration site identification (Wells et al., 2020). Lastly, a recent work also makes use of UMI-tag to achieve to quantify small somatic variant calling from ctDNA sequencing data (Hu et al., 2022). Still, there is to date no systematic comparative evaluation of a UMI-based system against fragment-based methods for abundance quantification in the context of insertion site retrieval using a set of controlled “real-world” scenarios. We here present a novel computational pipeline for integration site (IS) retrieval called IS-Seq which is capable of detecting IS at high resolution while allowing a user to leverage different tools for IS quantification including UMI abundance. We describe the overall design of the IS-Seq pipeline as well as the oligos used in our wet-lab protocol for IS retrieval, including a unique UMI configuration to specifically address sequencing errors. We then present the results of our analyses comparing the performance of IS-Seq and INSPIIRED for IS retrieval and quantification, as well as an evaluation of the different abundance estimation methods embedded in IS-Seq.

## RESULTS

### Wet-lab protocol and oligos design

The IS-Seq computational pipeline is tailored for analyzing the sequencing output of a customized ISA protocol previously described (Leonardelli et al., 2016; Biasco et al., 2018). Our wet-lab procedure resembles closely the latest iteration of insertion site retrieval techniques utilized by others in the field and is built on the wet-lab method originally designed by Schmidt et al (Gabriel et al., 2009) (Supplementary Fig.1A). Briefly, our ISA protocol is based on the fragmentation of gDNA by sonication, followed by ligation of a linker cassette (LC) with a known sequence, and by the exponential PCR amplification of the resulting fragments using vector LTR-specific and LC-specific primers. A final PCR step is carried out with primers that introduce into each sample Illumina adaptor sequences and a known sample index sequence: this allows pooling multiple samples for NGS on the Illumina platform and to demultiplex the results using an index-based assignment of the sequencing reads to each original sample. The oligos used in this protocol for the detection of lentiviral integration sites are shown in Supplementary Table 1. It is beyond the scope of this manuscript to describe the entirety of this procedure in detail, however, to facilitate the interpretation of the IS-Seq readouts, we will focus here on the custom design of our linker cassette and of the custom Illumina adapters (Fusion primers, Supplementary Tables 2,3).

The linker cassette used in our method carries a Unique Molecular Identifier (UMI) designed as shown in Supplementary Fig.1B, where two random 6nt stretches flank a 6nt “anchor” sequence (5’-GTAAGG-3). Differently from UMIs used in other protocols, the addition of an anchor sequence in our design allows us to assess the quality of sequencing specifically in the UMI region. For our analytical purposes we can therefore discard sequences that have nucleotide/s mismatches in this area under the assumption that such sequencing errors might have extended to the 6nt random sequences at the 5’ and 3’of the anchor therefore affecting the UMI diversity readout. Downstream the UMI the LC2 sequence is the one that directly attach to the genomic fragments while, upstream the UMI, the LC1 sequence is designed to host the binding of reverse primers for PCR amplification of vector genome junctions and for fusion PCR with custom Illumina adapters (fusion primers). The fusion primers used in our pipeline are designed according to Illumina protocol specification carrying at their 5’ end either the P5-Read1(for the LTR fusion primers) or P7-Read2 (for the LC fusion primers) Illumina sequences. To facilitate cluster recognition upon sequencing these custom adapters have an additional 12nt random sequence (5’-NNNNNNNNNNN-3’ in Supplementary Tables 2,3) immediately downstream Read1 or Read2. At the immediate 3’ end of this 12nt random stretch, 8nt sample barcodes are placed on both LTR and LC fusion primers, the combination of which is used for sample-specific tagging and demultiplexing by the computational pipeline as described below. Lastly, the 20nt segments at the 3’ end of the fusion primers, are designed to bind specifically to the LTR or LC sequence.

### Overview of the IS-Seq computational pipeline

The IS-Seq pipeline is designed to convert raw Illumina sequencing BCL files into a final table containing information of the genomic localization of integration sites (including annotation of the nearest gene) and their relative abundance per sample. To this aim, a series of steps are implemented based on quality filters, retrieval of fragment lengths and UMIs, mapping to the reference genome and annotating the results, as schematized in Fig.1,2. The first readout of the pipeline always expresses IS abundance in terms of absolute sequencing reads. The pipeline can then be run using the argument -a “fragment” or -a “umi” to obtain abundance results based on fragment lengths or UMI counts (see Material and Methods). The IS-Seq pipeline utilizes Python Threading (https://docs.python.org/3/library/threading.html) to analyze different samples concomitantly so that the average computational time for processing a MiSeq run is 4 hours. Given the length of this pipeline, we added “checkpoints” that, if needed, allow resuming the run from intermediate steps without restarting from the beginning. The pipeline works using as input BCL files in combination with library information contained in the “association file”. This *.csv file contains a datasheet that must be prepared according to the instructions reported in the GitHub page. Importantly, the “PT-Transduction-ID” field in this file is used by the pipeline to retrospectively identify independent group of samples (such as the ones deriving from different patients or transduction experiments) among which we don’t expect to detect shared identical insertion sites.

**Figure 1:**
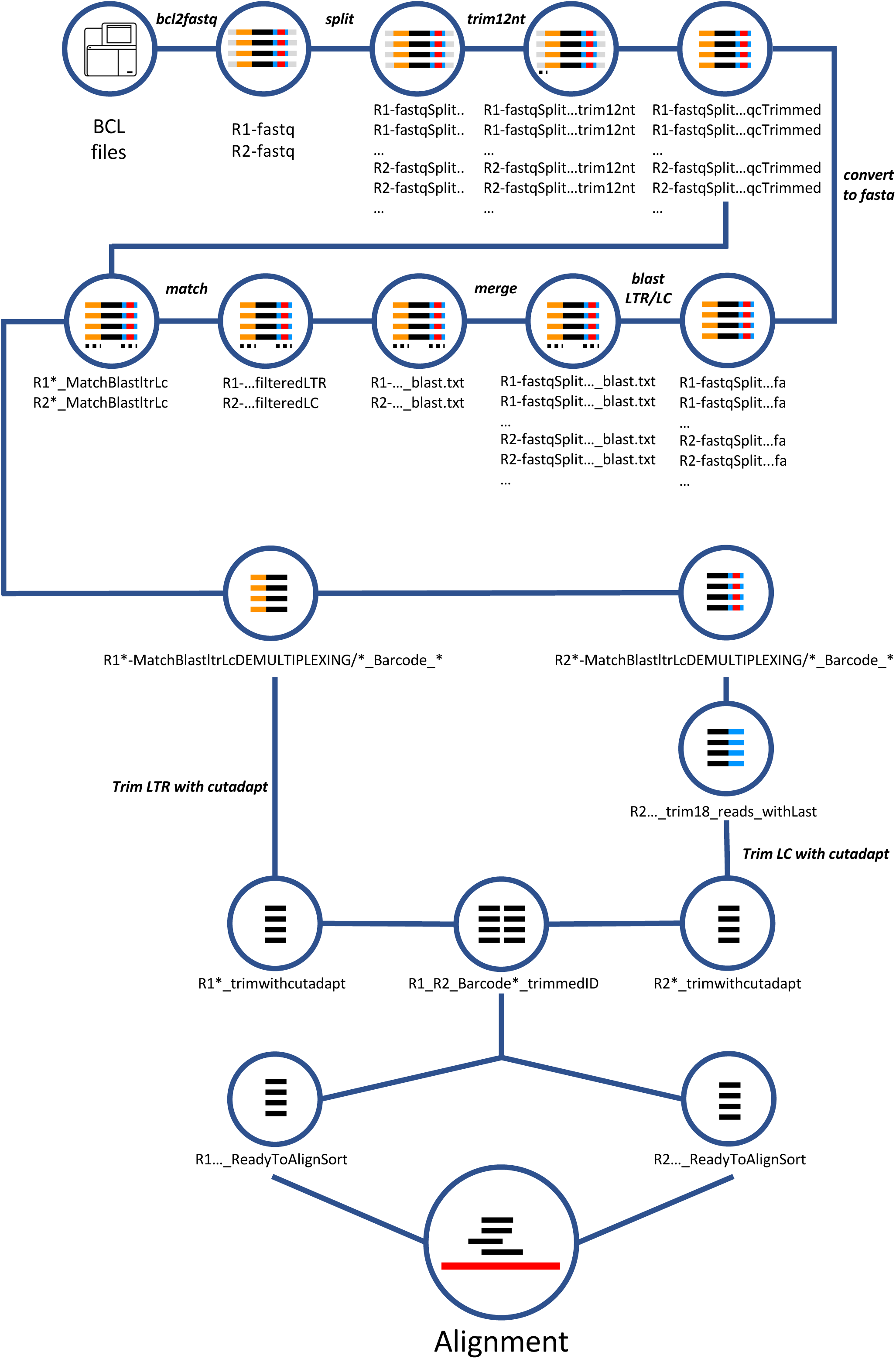
Schematic representation of the computational steps of the IS-seq pipeline from processing raw BCL Illumina files to right before the alignment step (LTR = Long Terminal Repeat, LC = Linker Cassette)

The first part of the workflow consists in obtaining R1 and R2 FASTQ files carrying the genomic fragments ready for alignment against the reference genome (Fig.1). To start, the BCL files need to be converted to FASTQ files using the *bcl2fastq* package (Illumina, 2018). This step is run independently and separately from the rest of the pipeline (see Material and Methods). Once the FASTQ files are obtained the IS-Seq pipeline can be launched with a single command as described in the GitHub page and the rest of the steps will run automatically. Firstly, the FASTQ files are split for better computational handling and the pipeline proceeds then to identify and trim the 12 random nucleotides in our fusion primers. After this step the pipeline converts FASTQ files into FASTA file and look for the presence of the long terminal repeat (LTR) vector sequence on R1 and the linker cassette (LC) sequence on R2 using *Blast* against sequence files of reference (these files can be customized according to needs and applications). Our pipeline is designed with a conservative approach regarding the quality of the LTR match, because of the importance of this segment for appropriately identifying vector-genome junctions. Therefore, we discard all sequences that do not have at least a 90% match with the LTR and LC sequences. At this stage, the pipeline looks for sample barcodes and performs demultiplexing of LTR and LC barcodes independently. The LTR sequence in R1 is then trimmed with *cutadapt* and the remaining sequences are stored as FASTQ files. Likewise, LC is trimmed from R2, but these sequences are further processed for UMI recognition. The IS-Seq pipeline first identifies and checks for perfect match with the anchor sequence (“ancora” in the script), then looks at the flanking 6nts upstream and downstream the anchor. Lastly, the pipeline trims the 18nt sequence and stores the information for abundance estimation via UMI quantification (see dedicated Results section). At the end of this process, sequence IDs in R2 are ready to be matched with the ones in R1. The R1 and R2 FASTQ files with matching IDs are then selected for going through the alignment step.

Alignment is performed using the Burrows-Wheeler Aligner (BWA) algorithm (Li and Durbin, 2009) and the output are *.sam files for each R1/R2 barcode combination which are converted to .bam files to undergo additional filters (Fig.2). Firstly, the IS-Seq pipeline identifies the reads that are aligned to host genome repetitive regions and filters the reads with low mapping quality by MAPping Quality (MAPQ) score. After this step, the IS-Seq pipeline outputs the reads that are not aligned to repetitive regions and combines these reads with the reads found in repetitive regions but having high mapping quality, into the R1_R2*_allFilter.bam file. The IS-Seq pipeline then filters all alignments in R1_R2*_allFilter.bam file that do not have a perfect match of the 3nt at the beginning of the genomic fragment immediately downstream where the LTR originally was before trimming. This is to ensure that only bona-fide vector-genome junctions are carried over for identifying the insertion site of the vector. The “*nonSupplementary.bam” files are then selected for final parsing of the results. At this stage our pipeline is designed to output the alignments without further filters (filterNo) and the ones that survive an additional quality filter based on selecting only the alignments with at least a 60bp long genomic match (filter60) of the sequence originally flanking the LTR. This is again to warrant a conservative quality assessment of vector-genome junctions used for insertion sites identification.

**Figure 2:**
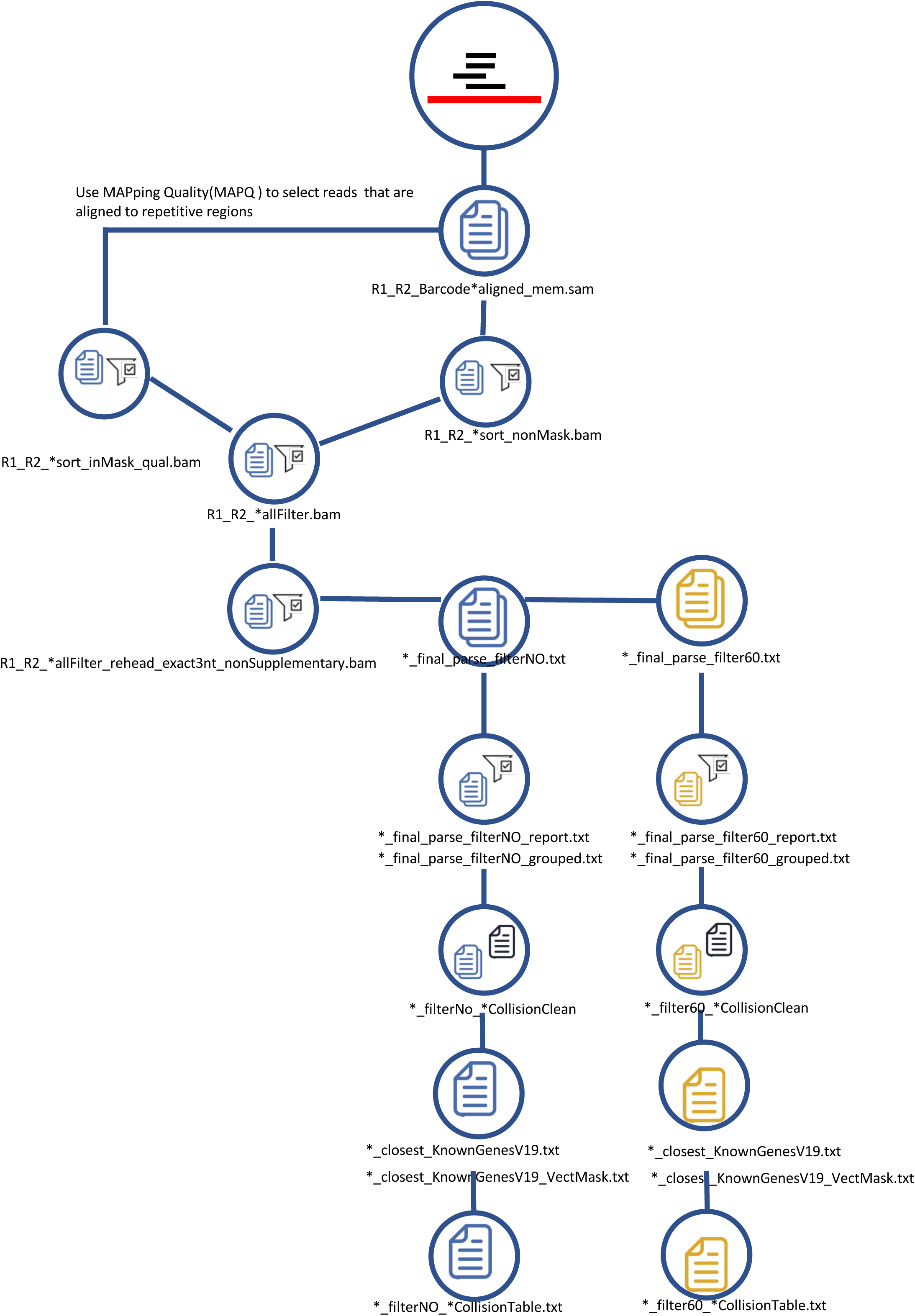
Schematic representation of the alignment procedure embedded in the IS-seq pipeline and the filtering steps generating the final output tables.

The IS-Seq pipeline calls insertion sites falling within a 7 bp window as identical insertions. To this aim, the pipeline selects the alignment position associated with the highest abundance and then merges the insertions in the 7bp region into a single entry with abundance corresponding to the sum of all abundances. This 7bp window was established based on tests on simulated and “real-world” datasets, where we measured the variance of total integration sites obtained using different merging windows from 0bp to 50bp (Supplementary Fig.2). We observed, that when window size changes from 0bp to 3bp the number of integration sites decreased progressively due to the merging of integration sites falling within this window. However, from windows of 3bp to 7bp, the pipeline output converged rapidly to stable numbers of IS which are therefore likely to represent true independent events. Therefore, we used the 7bp window as a conservative cutoff to merge proximal integration sites under the reasonable assumption that everything closer than 7bp most likely represent noise/artifacts or local fraying events which belong to a single IS event. Similar results and conclusions have been reported in another publication (Spinozzi et al., 2017). Lastly, samples are grouped according to the PT-Transduction-ID of origin as reported in the association file. At this stage rare identical genomic locations (“collisions”) identified across independent PT-Transduction-ID groups and defined again as IS falling within a 7bp window from each other, are re-assigned based on a “10-fold abundance rule” (Aiuti et al., 2013; Biffi et al., 2013; Biasco et al., 2015) to the PT-Transduction-ID carrying a 10-fold higher normalized abundance than the rest of the groups. If an insertion is shared by two or more independent PT-Transduction-IDs with similar abundance (below the 10-fold threshold), such integration site is not assigned to any of the groups and is discarded. The 10-fold abundance rule idea is borrowed from RNA-Seq based Differential Gene Expression (DGE) analysis. Usually, in DGE, log2FC >=1.5 is used to define differential genes between two condition. That means FC should be greater or equal to 2.83. The number of reads belonging to integration sites across genome follow a similar negative binomial distribution as the number of reads across all genes in a gene expression study. Therefore, we decided to introduce this filter but with a more conservative approach using FC between two samples greater than 10. Lastly, insertion sites are annotated and filtered for positions mapping in the vector genome (“VectMask”). The “*CollisionTable.txt” files are the final output of the pipeline, consisting of a data frame with insertions sites and raw abundance by sample, to be used for further downstream analyses.

### Datasets used for testing IS-Seq performance

To test the performance of IS-Seq for insertion sites detection and quantification, we performed a series of comparative analyses against INSPIIRED, as the most recent publicly available suite for insertion sites detection. The results described in the following sections have been generated using *in silico* and *in vitro* datasets. Firstly, based on the design of the wet-lab protocol for IS-Seq, we performed extensive simulation studies by generating different simulated data sets with 1, 5, 100 or 1000 integration sites as ground truth, and tested the ability of IS-Seq to detect the ground truth in absence/presence of mutations and background genomic noise. For the dataset containing 100 IS, we also generated simulations with one ground-truth IS having 100%, 99%, 75%, 50%, 10%, 1%, 0% abundance with respect to other IS, and testes *in silico* conditions with different levels of PCR duplicates or different UMI entropies. More details regarding these simulations are reported in the Material and Methods section of the manuscript. We then analyzed a test dataset from INSPIIRED (available at https://github.com/BushmanLab/intSiteCaller) and sequencing data from 2 experimental conditions with HL60 cell line clones carrying known and traceable insertion sites of a Self-Inactivating Lentiviral Vector (SIN-LV) carrying the GFP marker gene under a PGK promoter. Specifically, the first analysis was performed on 9 SIN-LV transduced clones generated by single cell sorting and 27 days of expansion, carrying either 1,2,3 or 5 lentiviral copies per cell, as validated by ddPCR for vector copy number (VCN). The second set of analyses were performed instead on serial dilutions of gDNA from one clone (CL6) with VCN=1 into a polyclonal transduced HL60 population. All raw outputs of the different pipeline iterations can be found in the supplementary information section.

### Testing IS-Seq performance on insertion site datasets from *in silico* simulations and INSPIIRED

To assess the detection and quantification potential of our IS-Seq computational pipeline, we first generated multiple simulated datasets as described in the material and methods section. Simulation 1 was based on extracting in silico a 3kb fragment from the hg38 human genome and generating a simulated insertion sites associated to 2,924 reads in the chr19, 49461738 position appending LTR and LC sequences at the 5’ and 3’ ends respectively according to the expected R1 and R2 outputs of our wet-lab/sequencing procedure. As shown in supplementary table 4, the IS-Seq pipeline returned a perfect match with the input FASTQ file (2,924 reads in the chr19 49461738 position). In simulation 2 we artificially introduced mutations in the first 50bp after the LTR in 10% of the 2,924 input reads (The mutation rates of the mutated 50bp sequence ranging from 0.4651163 to 0.9230769) which we expected to be discarded by IS-Seq upon filtering and we obtained again the expected results with 2,632 reads mapped in the chr19 49461738 position. For simulation 3 we also added genomic background noise in the form of 17,274,461 random reads. IS-Seq returned the same results with 2,632 reads mapped in the chr19 49461738 position without detecting any false positives. In simulation 4 we included 4 additional insertions sites as input material where 75% of the reads (n=3006) belonged to the simulated insertion in the chr19 49461738 position while 25% to 4 insertions in other genomic positions, 2 of which expected to map in repetitive regions (supplementary table 4). We also again added 17,274,461 random reads as genomic background noise. We then run IS-Seq and obtained the results shown in supplementary table 4. IS-Seq correctly identified and annotated 3006 reads in the chr19 49461738 position (75% relative abundance), 251 reads in the chr19 40461720 position and 251 reads in the chr19 2036173 position. As expected, reads belonging to remainder two additional insertions were split over 2 and 6 mapping positions respectively as a result of a non-unique mapping without any loss of reads. Again, no additional insertion was detected from 17,274,461 random genomic reads introduced as background noise. In simulation 5 and 6, we set to produce integration sites across the whole human genome assigning the number of integration sites proportionally to each chromosome based on its relative size, generating 100 or 1000 random integration sites respectively (including the one in the chr19 49461738 position). We then evaluated the performance of IS-Seq on this simulated data and obtained high-performance metrics in terms of Precision-Recall (PR), Receiver Operating Characteristic (ROC) curves as well as of the proximity between estimated and expected abundance (Supplementary Fig. 9). Altogether these in silico simulations showed that IS-Seq had the potential to accurately detect insertion sites mapping in unique chromosomal locations, to properly filter mutated sequences and background noise as well as to preserve and collect insertion sites mapping in repetitive regions.

Given the limitations of inherently artificial *in silico* simulations in reproducing “real world” scenarios, we next conducted a series of additional tests on data generated from *in vitro* experiments. To this aim, we firstly analyzed the test dataset provided on the INSPIIRED GitHub webpage to directly compare IS-Seq performance against INSPIIRED in absence of potential biases associated with the different wet-lab protocol and sequencing conditions we use for IS retrieval. This dataset is composed of 3 FASTQ files from 8 vector-positive clones or a combination thereof (pool), each of which analyzed in 4 replicates for a total of 32 samples. Since the format of the INSPIIRED dataset is different from that used as input by the IS-seq pipeline, we developed a procedure to reformat these data to an input format suitable for IS-Seq and adjusted the parameters accordingly. The details of this procedure are described in the Material and Methods section. Despite the dataset being the result of a different wet-lab IS retrieval protocol specifically designed for INSPIIRED, with this adaptation, IS-Seq was still able to identify 7 out of the 9 IS retrieved using INSPIIRED, the two missing being background integrations constituting 0.08% of the total fragment counts each (Fig.3). To quantify the degree of concordance between these two computational platforms, we artificially appended the 2 missing IS to the IS-Seq results and assigned an abundance of 0 to both. We then generated two 9*32 matrixes, one for each computational pipeline, using in both cases as measure of IS abundance the Maximum Likelihood Estimation (MLE) of fragment lengths (the SonicAbundance method in the INSPIIRED pipeline). To calculate the distance between the 2 matrixes, we applied a multivariate statistical approach using the Procrustes analysis (Oksanen et al., 2020). Using this method, we observed that the distance between the integration profile of INSPIIRED and the one obtained with IS-Seq was very low (0.000464197). To estimate the likelihood of such an outcome, we subsampled 500 random matrixes from the 9*32 IS-Seq matrix and calculated the Procrustes distance between the INSPIIRED profile and each of these random samples. We analyzed the distribution of the Procrustes distance for these 500 samples to estimate how often this distance was equal or less than 0.000464197, showing that in all cases the distance was higher than this observed value (Supplementary Fig. 2). The result of this analysis allowed us to estimate as statistically negligible the probability that the distance between INSPIIRED and IS-Seq results would equal 0.000464197, therefore supporting the notion that we obtained a high level of concordance between the results of these two pipelines. We also compared the abundance calculated using IS-Seq or INSPIIRED to the expected abundance of each clone in this dataset by calculating the Mean Square Error (MSE) and observed IS-Seq results were as close to the true values as the one obtained with INSPIIRED (Table 1). These results suggest that IS-Seq can be adapted to reliably analyze data generated with the INSPIIRED protocol, a first indirect validation of our pipeline as a reliable method for IS identification and abundance estimation

**Figure 3:**
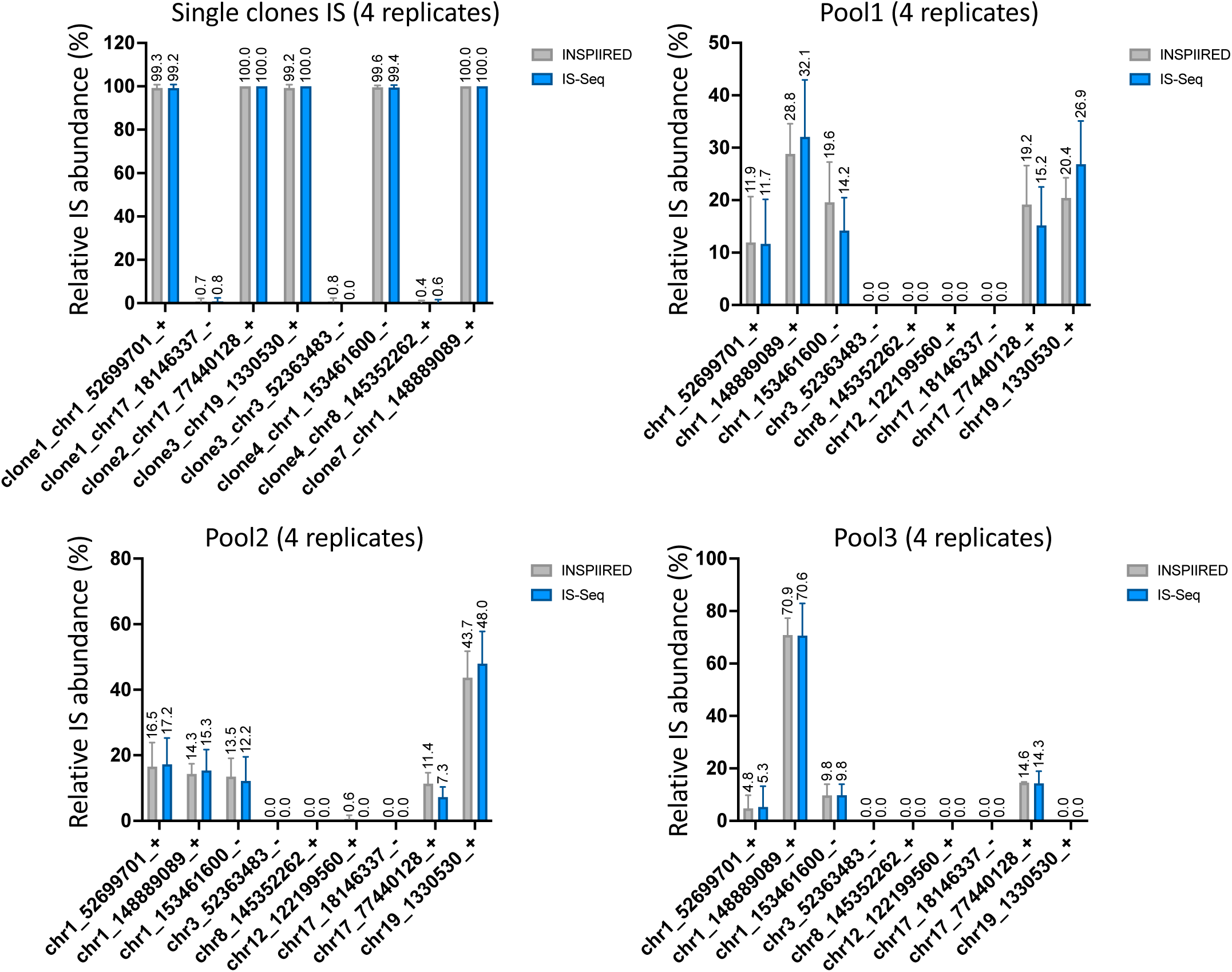
Relative abundance of insertion sites (IS) measured in single cell clones and pools from the publicly available INSPIIRED dataset using either the INSPIIRED (grey bars) or the IS-Seq (blue bars) pipelines. Mean and standard errors are reported for the 4 replicates relative to each sample. (MLE = Maximum Likelihood Estimate of fragment lengths).

**Table 1:** Mean square error (MSE) calculated between Is-Seq Fragment_MLE and INSPIIRED Fragment_MLE with known relative abundance of IS of all clones on INSPIIRED data set(https://github.com/BushmanLab/intSiteCaller/blob/master/testCases/intSiteValidation/t ruth.csv). The average MSE for all the results of each pipeline iteration is reported in the last row of the table (Fragment_MLE = Maximum Likelihood Estimation of Fragment Lengths)

| Sample | MSE_INSPIRED | MSE_ISSeq |
| --- | --- | --- |
| Clone1(1 IS) | 0.49 | 0.64 |
| Clone2(1 IS) | 0 | 0 |
| Clone3(1 IS) | 0.64 | 0 |
| Clone4(1 IS) | 0.16 | 0.36 |
| Clone7(1 IS) | 0 | 0 |
| Pool1(5 ISs) | 28.802 | 63.918 |
| Pool2(5 ISs) | 6.166666667 | 10.01 |
| Pool3(5 ISs) | 30.53 | 29.116 |
| Average MSE | 8.348583333 | 13.0055 |

### Testing IS-Seq performance to identify and quantify IS in single cell clones

To establish the ability of our wet-lab protocol combined with the IS-Seq computational pipeline to accurately identify IS and to estimate their relative abundance, we firstly analyzed cell clones with known and uniform vector copy number (VCN). To this aim we generated single-cell HL60 clones and assessed their copy number by ddPCR (digital droplet PCR) as described in the Material and Methods section. We obtained 9 stable and uniform cell clones, 1 of which with a VCN=1, 6 with VCN =2, 1 with VCN =3 and 1 with VCN=5. We then applied IS-Seq to these datasets to establish whether we could detect the expected number of insertion sites in each clone with the expected relative abundance (e.g. from the clone with VCN=5 we should have been able to identify 5 IS with approximately 20% relative abundance each). As performance comparator we used INSPIIRED providing as input R1 and R2 sequencing data preprocessed with IS-Seq before alignment (see Material and Methods). In Fig. 4 and Supplementary Fig. 3 we summarized the results of our analyses based again on the use of MLE of fragment lengths quantifications for both pipelines. In both cases we were able to detect the main and most abundant vector insertion sites belonging to each clone in the expected numbers accompanied by a negligible fraction of background insertions with very low abundance (equivalent to a combined average of 1.5% for IS-Seq results), which in this context can be ascribed to contaminants, PCR artifacts or sequencing noise. We again estimated the Procrustes distance between INSPIIRED and IS-Seq which we measured as being 0.265555, higher than what was observed in the previous test on the INSPIIRED dataset. However, when we measured the correlation coefficient between IS-Seq and INSPIIRED results for each clone, we could show that this was very high ranging from (0.97 to 1) in all clones except for CL6 where it was 0.18 (Supplementary Table 5). We identified this particular result as the one most affecting the overall Procrustes distance between the 2 pipelines. Indeed, we noticed that, when analyzing this specific clone (with VCN=1) with INSPIIRED we lost most reads belonging to the main IS. Inspecting these results more closely we identified that the reason was the 95% sequence similarity threshold (SST) used upon alignment with BLAT as per INSPIIRED default configuration. Indeed, by artificially lowering this parameter we observed a substantial increase in the retrieval of sequence reads belonging to the most abundant IS of CL6, as addressed below. Once established that the results of the two pipelines indeed have a high level of concordance, we went on measuring the distance between the theoretical and observed abundance of the most abundant IS in each of these clones for IS-seq and INSPIIRED. Because of the adjustments required for the INSPIIRED pipeline to detect the IS in CL6, for a fair comparison between the two pipelines we estimated the distance between expected and observed results in INSPIIRED using sequence similarity thresholds of either 95%, 80% or 0% (Fig. 4 and Supplementary Fig. 3). The results of this analysis are reported in Table 2, where we compared the MSE for each clone using each of the computational pipelines. The average MSE of IS-Seq was 138.6944 while for INSPIIRED was 312.0519, ranging from 257.6178 to 364.8111 based on the use of the 3 sequence similarity thresholds mentioned above.

**Figure 4:**
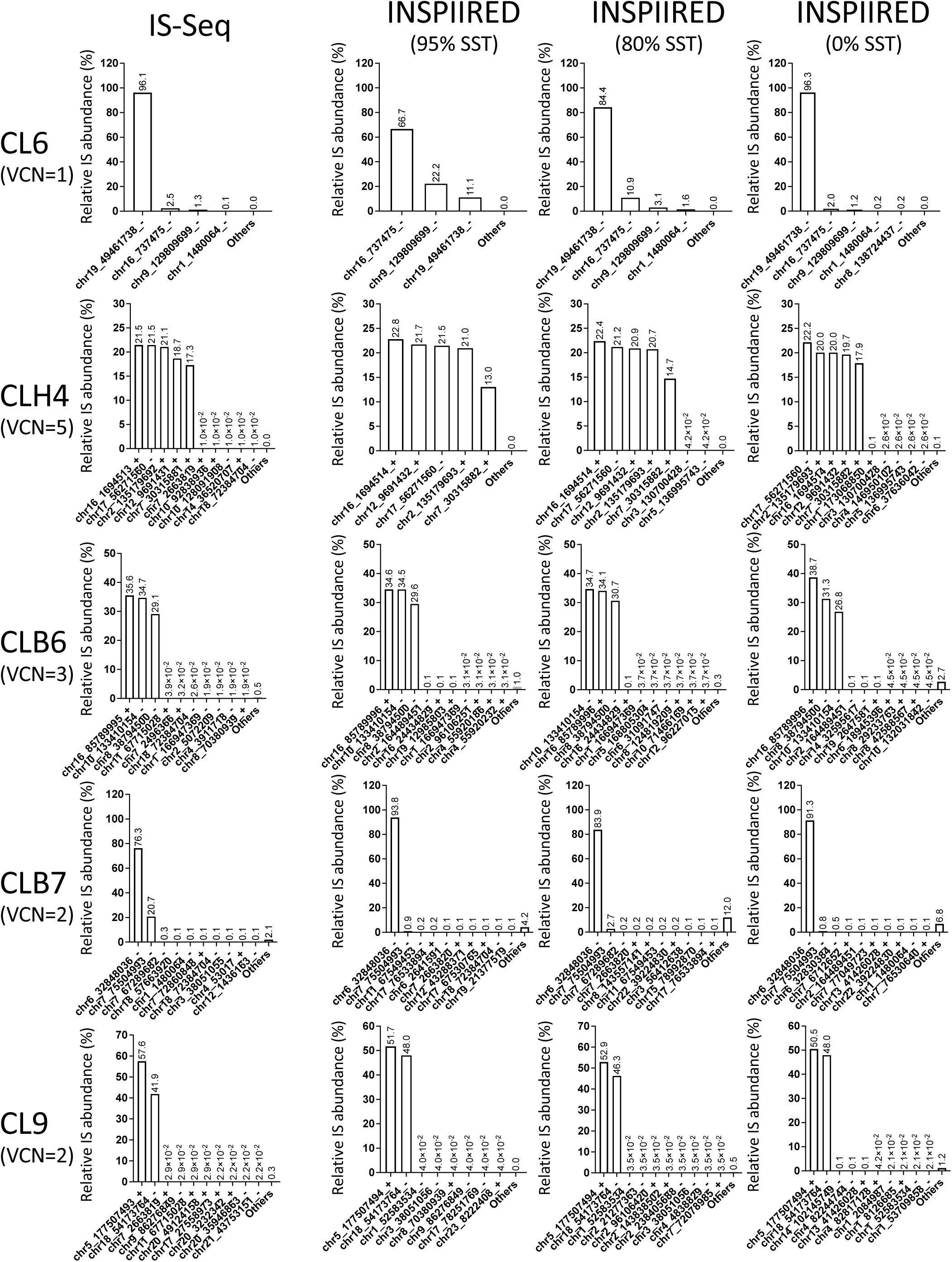
Relative abundance of insertion sites (IS) detected in single cell clones using different pipelines and sequence similarity thresholds (SST) based on the Maximum Likelihood Estimate of fragment lengths. (VCN = vector copy number) (results relative to the test of the clones can be found in Supplementary Fig. 4)

**Table 2:** Mean square errors (MSE) calculated between the results of each pipeline iteration and the expected relative abundance of the most abundant IS (main IS) detected for each clone (see Fig.3 and Supplementary Fig.3 for the relative abundance results). The average MSE for all the results of each pipeline iteration is reported in the second last row of the table, while the global MSE calculated for IS-Seq or INSPIIRED is shown in the last row. (MLE = Maximum Likelihood Estimation, SST = Sequence Similarity Threshold)

| Sample | VCN | Expected<br>(% of<br>main IS) | Is-Seq<br>(Fragment_MLE)<br>(2326) | INSPIRED<br>(Fragment_MLE) |  |  |
| --- | --- | --- | --- | --- | --- | --- |
|  |  |  |  | 95%<br>(513) | 80%<br>(621) | 0%<br>(1314) |
| CL6 | 1 | 100 | 16.48 | 1111.09 | 244.14 | 14.00 |
| CLH4 | 5 | 5*20 | 2.90 | 12.43 | 7.23 | 1.85 |
| MOI30CLB6 | 3 | 3*33.33 | 8.30 | 3.29 | 5.76 | 25.03 |
| MOI30CLB7 | 2 | 2*50 | 722.53 | 1881.30 | 1691.12 | 2066.29 |
| MOI30CLC9 | 2 | 2*50 | 60.70 | 3.53 | 11.02 | 2.19 |
| MOI30CLE1 | 2 | 2*50 | 12.24 | 56.40 | 60.45 | 74.38 |
| MOI50CLB9 | 2 | 2*50 | 5.89 | 19.11 | 23.39 | 88.00 |
| MOI50CLE6 | 2 | 2*50 | 28.39 | 15.26 | 36.38 | 218.71 |
| MOI50CLH6 | 2 | 2*50 | 390.82 | 180.89 | 239.07 | 333.09 |
| Average MSE |  |  | 138.6944 | 364.8111 | 257.6178 | 313.7267 |
| Global MSE |  |  | 138.6944 | 312.0519 |  |  |

Notably, we observed that despite the threshold adjustments, the INSPIIRED pipeline was never able to detect with high efficiency the second most abundant integration of the CLB7 clone with VCN=2, which was instead called as IS in the chr7, 75504993 unique non-repetitive genome position with 20.7% relative abundance using IS-Seq, suggesting that the BLAT mapping protocol, used by INSPIIRED, might have some inherent limitations when dealing with insertions in the genome of immortalized cell clones (see the Discussion section for more details). To assess whether INSPIIRED classified reads belonging to this insertion as IS in repetitive elements, we used the “multihits” clustering procedure of INSPIIRED. We first appended the unique hits with sequence clusters of IS in repetitive elements (Supplementary Figure10) generated with this function at different sequence similarity thresholds. Looking at these results the top abundant insertion in CLB7 was now a sequence cluster in a repetitive element, differently from both the first (chr6_32848036) and the second (chr7_75504993) most abundant IS identified with IS-Seq in Fig.4. By simply adding reads mapping in repetitive elements to the one mapping in unique IS without clustering them by sequence similarity, we recovered more reads belonging to the second most abundant IS in chr7, 75504993 (Supplementary Figure11), but, in the process, the background noise of reads assigned to other positions also increased to levels that made the results completely detached from the ground truth. Most importantly, when running the multihits clustering procedure on other two clones, CLH6 and CL6, whose IS were validated as 2 and 1 unique hits respectively using both INSPIIRED and IS-seq, we observed that this procedure generated additional repetitive sequence clusters with high abundance on top of the IS mapping in unique positions previously identified (Supplementary Figure 10). This led us to the conclusion that the multihits clustering procedure, did not allow detecting with higher efficiency the second IS of the CLB7 clone, while instead increased the background noise impairing our ability to assess real-world ground truth IS content in this and other clones. Based on these findings, we advocate that the results of the “multihits” clustering procedure of INSPIIRED should be used with caution in contexts where there is no formal way to assess *a priori* the IS content of a given specimen such as when analyzing clinical samples.

Overall, these results underscore that our wet-lab protocol combined with IS-Seq is at least as (if not more) accurate and efficient than INSPIIRED in the identification and quantification of IS from clones with known VCN.

### Testing IS-Seq performance to quantify clone-specific IS upon dilution into polyclonal populations

In a clinical scenario where IS analysis is requested, samples would contain individual clones with dominant insertion sites mixed with a polyclonal population of cells with a diversity of viral integrations. To reproduce such a scenario in a controlled laboratory experiment, we generated serial dilutions of CL6 genomic DNA (gDNA) into a polyclonal population of bulk HL60 cells transduced with the same lentiviral vector construct as shown in Fig. 5A. The goal of this experiment was to test the performance of IS-Seq on quantifying the relative abundance of the CL6 insertion site and, secondarily, to test the capacity of our pipeline to detect the other insertions contained in the polyclonal population with a resolution comparable to INSPIIRED. To this aim we performed 3 independent dilution experiments where we applied our wet-lab protocol for insertion sites retrieval to either 250ng (250 and 250rep in Table 3) or 500ng (500 in Table 3) of gDNA as starting material. We then sequenced the relative libraries in 2 independent sequencing runs (250+500 in the first run and 250rep alone in the second run). To identify the insertion sites, we again used either an adaptation of INSPIIRED or our IS-Seq pipeline and quantified the insertional abundance by means of MLE of fragment lengths. Because of the previous observation relative to the inefficiency of the default mapping parameters of INSPIIRED to detect the CL6 insertion, we run this pipeline using either 95%, 80% and 0% sequence similarity thresholds. The results are shown in Fig. 5B, where we plot the relative abundance of CL6 measured with either IS-Seq or INSPIIRED, and in Table 3 where we reported also the average MSE calculated against the expected values for each dilution experiment and the overall MSE per pipeline iteration. When comparing the results of IS-Seq and INSPIIRED against the expected values we observed that at higher dilutions (less than 10% of CL6 insertion site content) as well as when only CL6 clone gDNA was used as input material (100% CL6 insertion site content) the two pipelines performed in a relatively comparable fashion. However, when we analyzed the 10%, 50% and 75% dilution series, we observed that IS-Seq clearly outperformed INSPIIRED in terms of accuracy of clone abundance estimation regardless of the sequence similarity thresholds used for the latter. As a result, the average MSE value of IS-Seq (123.5367) was lower than any of the 3 INSPIIRED iterations (2897.03 using the 95% threshold, 922.66 using the 80% threshold and 336.8333 using the 0% threshold). Regarding the number of insertions detected in the polyclonal population, we could show that IS-Seq and INSPIIRED had comparable efficiency to detect insertions sites other than CL6 at all dilutions. Specifically, when comparing the 0% dilution data point (only HL60 polyclonal population used as input) using 500ng as starting materials, we could observe that the overlap of IS detected in the two pipelines (using INSPIIRED with 0% sequence similarity threshold) was of 855 IS, with 140 IS and 155 IS detected respectively by the INSPIIRED or IS-Seq pipelines only (Supplementary Fig. 4). Overall, these data show that IS-Seq is at least as (if not more) efficient and accurate than INSPIIRED to estimate the contribution of a given clone in a mixed cell population as well as to identify and quantify viral integrations in a polyclonal population. However, this experiment highlighted some limitations of the IS abundance estimation system based on MLE of fragment lengths, particularly for what concerned the accuracy of the 75% and 50% clonal dilutions estimation. For this reason, we implemented in our IS-seq pipeline the UMI diversity abundance estimator and tested its performance in the same experimental conditions against the most widely used MLE of fragment lengths.

**Figure 5:**
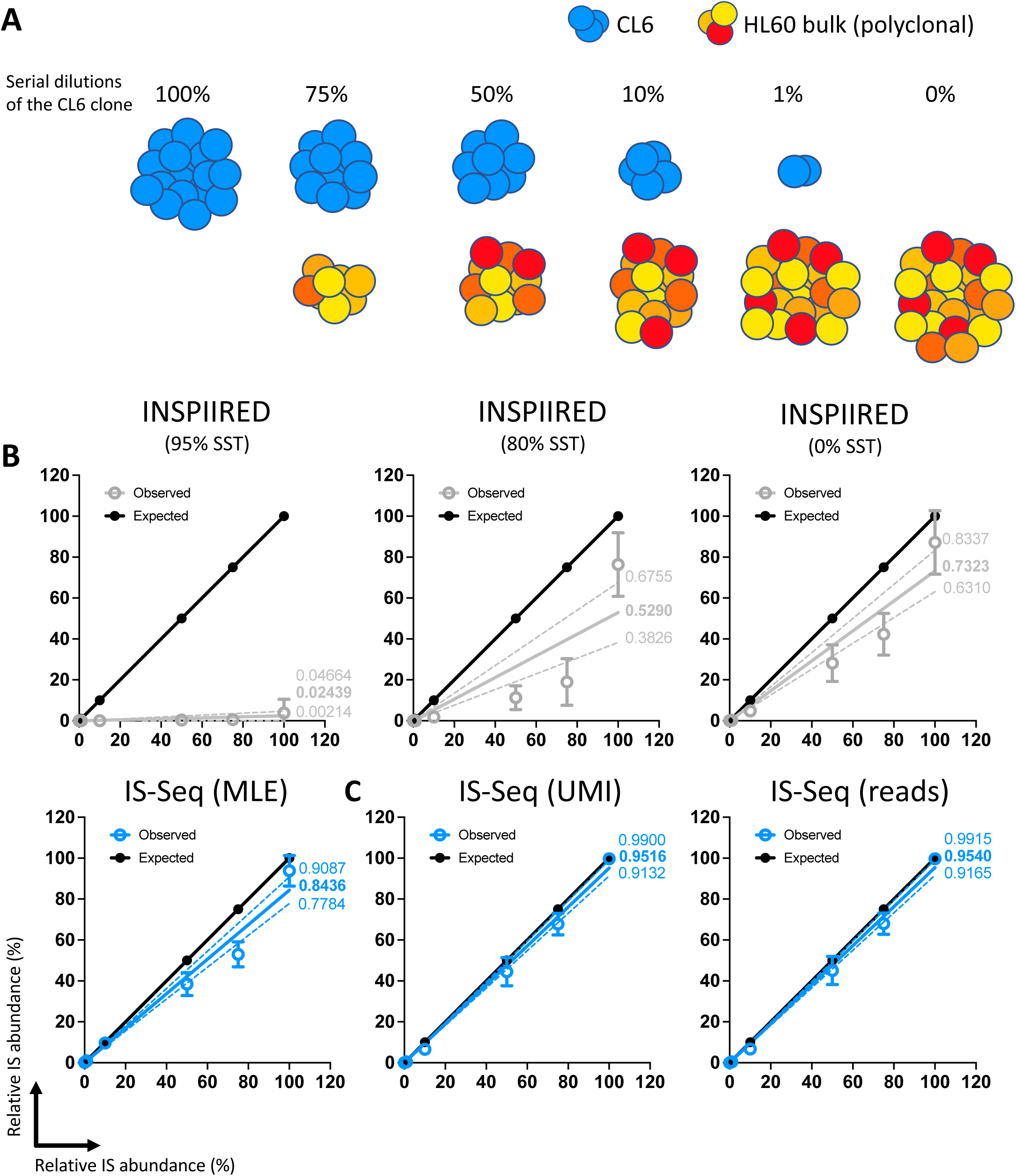
A) Schematic representation of the serial dilutions of CL6 (in blue) into the bulk polyclonal HL60 population (scale of oranges and reds). B) Linear regression of expected relative abundance values (in black) vs observed relative abundance values using either IS-Seq (in blue) or different iterations of the INSPIIRED pipeline (in grey). All abundance are measured with Maximum Likelihood Estimate of Fragment Lengths (MLE). Mean and errors are shown as dots and error bars. The full line show the linear correlation (slope value reported in bold) while dotted line the 95% confidence interval (min and max slope values reported in normal font). (SST = Sequence Similarity Threshold, IS = CL6 insertion site) C) Same analysis performed on IS-Seq results generated with the Unique Molecular Identifiers (UMI) or Sequence reads quantification methods.

**Figure 6:**
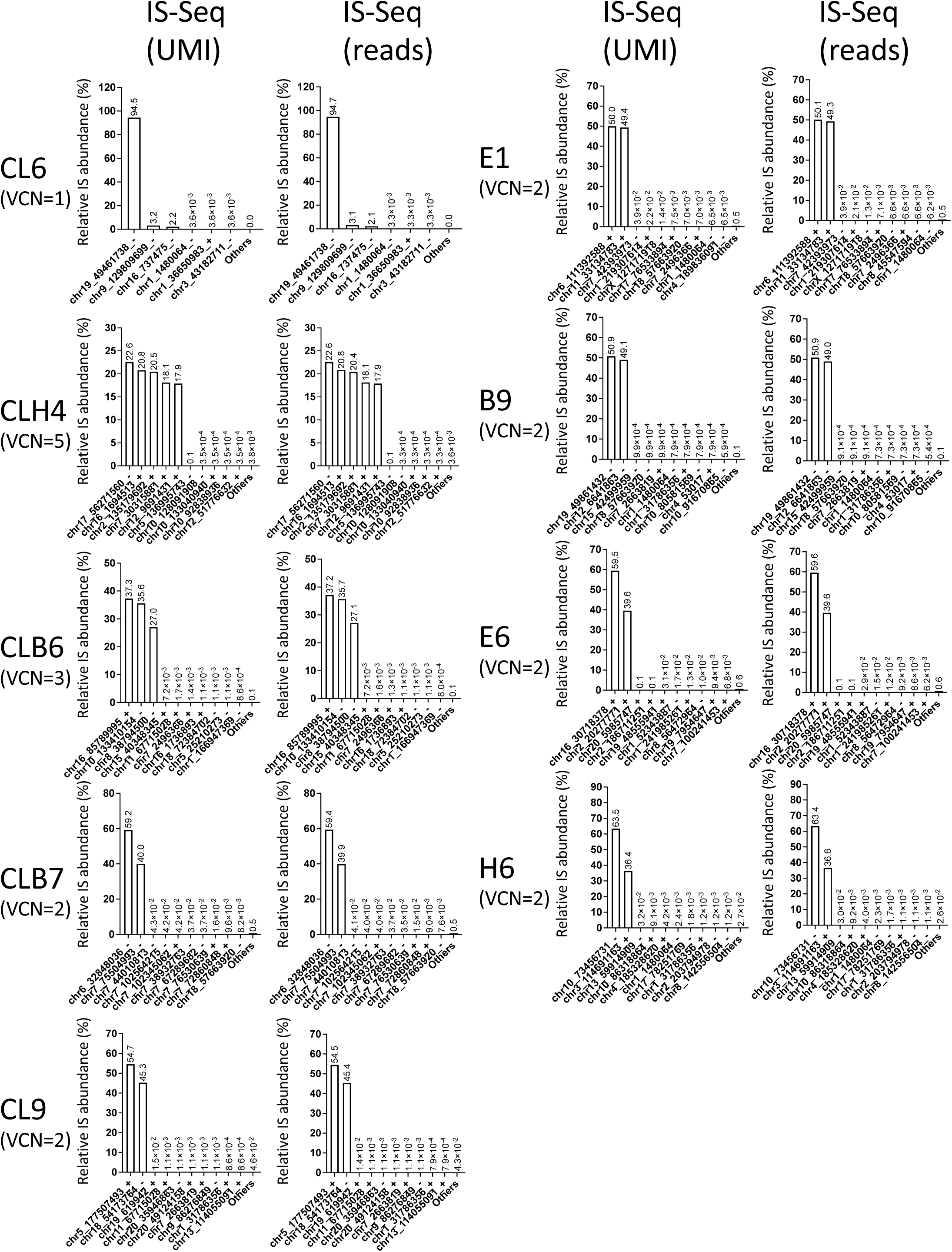
Relative abundance of insertion sites (IS) detected in single cell clones using IS-Seq based on either Unique Molecular Identifiers (UMI) or Sequencing reads (reads). (VCN = vector copy number)

**Table 3:** Expected and observed contribution of the CL6 insertion site upon serial dilution into a bulk HL60 transduced polyclonal population, as measured by IS-Seq and INSPIIRED. In the second last row is reported the Mean square error (MSE) between the results of each pipeline iteration and the expected relative abundance of the serial dilutions of CL6. The average MSE for all the results of each pipeline iteration is reported in the last row of the table. (MLE = Maximum Likelihood Estimation, SST = Sequence Similarity Threshold)

|  | IS-Seq<br>(Fragment_MLE) |  |  | INSPIRED<br>(Fragment_MLE) |  |  |  |  |  |  |  |  |
| --- | --- | --- | --- | --- | --- | --- | --- | --- | --- | --- | --- | --- |
|  |  |  |  | 95% SST |  |  | 80% SST |  |  | 0% SST |  |  |
| Expected (%) | 250<br>(n=1793) | 500<br>(n=3463) | 250rep<br>(n=2792) | 250<br>(n=1080) | 500<br>(n=2071) | 250rep<br>(n=1080) | 250<br>(n=1207) | 500<br>(n=2319) | 250rep<br>(n=1120) | 250<br>(n=1802) | 500<br>(n=3345) | 250rep<br>(n=2596) |
| 0 | 0.102 | 0.000 | 0.318 | 0.000 | 0 | 0 | 0.000 | 0.000 | 0.194 | 0.000 | 0.000 | 0.244 |
| 1 | 0.455 | 0.543 | 1.807 | 0.000 | 0 | 0 | 0.000 | 0.000 | 0 | 0.146 | 0.175 | 0.285 |
| 10 | 9.916 | 8.108 | 10.931 | 0.000 | 0 | 0 | 1.515 | 1.443 | 2.602 | 4.086 | 4.232 | 6.246 |
| 50 | 39.640 | 32.362 | 43.331 | 0.000 | 0 | 1.081 | 11.397 | 5.416 | 17.035 | 25.245 | 21.041 | 38.184 |
| 75 | 55.847 | 46.021 | 57.026 | 0.529 | 0 | 1.149 | 14.122 | 10.766 | 32 | 39.965 | 33.496 | 53.526 |
| 100 | 97.085 | 99.099 | 85.303 | 0.000 | 0 | 11.538 | 81.538 | 88.679 | 58.889 | 93.758 | 98.286 | 69.429 |
| MSE | 80.50 | 192.58 | 97.53 | 3024.49 | 3037.6 | 2628.9 | 935.03 | 1052.69 | 780.26 | 319.15 | 433.02 | 258.33 |
| Average MSE | 123.5367 |  |  | 2897.03 |  |  | 922.66 |  |  | 336.8333 |  |  |

### Testing the performance of IS abundance estimation using UMI diversity and comparison with MLE of fragment lengths

As reported above, the IS-Seq pipeline can retrieve and store information relative to the abundance of UMIs embedded in our linker cassette design (Fig. 1) and to use their diversity for IS abundance estimation. To test the performance of this quantification method in a controlled experiment, we first run IS-Seq to identify and quantify IS in the single cell clones above described, using the UMI abundance estimation option (see Material and Methods). The relative abundance of the insertion sites of each clone as measured by UMI diversity is shown in Fig 6 where we also report the abundance calculated using sequencing reads counts as a benchmark control. We then quantified the distance between observed and expected results and obtained the MSE values reported in Table 4. In the same table we also reported the results of the IS-Seq pipeline run on the same samples using the MLE of fragment length. The average MSE across all samples using the UMI-based abundance estimation was 50.15 which was substantially lower than the average MSE obtained using the MLE of fragment lengths (138.69). Notably, in this scenario, the MSE obtained using the Read-based abundance estimation (49.82) showed that this method also outperformed the MLE of fragment length and that it was in line with the accuracy of the UMI quantification system. We then applied the same approach to the serial dilution experiment described in the previous paragraph to assess the performance of the UMI-based vs MLE of fragment length using IS-Seq.

**Table 4:** Mean square errors (MSE) calculated between the results of IS-Seq using different quantification methods and the expected relative abundance of the most abundant IS (main IS) detected for each clone. The average MSE for all the results of each pipeline iteration is reported in the second last row of the table, while the global MSE calculated for each IS-Seq quantification method is shown in the last row. (MLE = Maximum Likelihood Estimation, UMI = Unique Molecular Identifiers)

| Sample Name | VCN | Is-Seq Reads<br>(n=2490) | Is-Seq UMI<br>(n=2490) | IS-Seq<br>Fragment_MLE<br>(n=2326) |
| --- | --- | --- | --- | --- |
| CL6 | 1 | 27.99 | 29.98 | 16.48 |
| CLH4 | 5 | 3.09 | 3.12 | 2.9 |
| MOI30CLB6 | 3 | 19.96 | 20.25 | 8.3 |
| MOI30CLB7 | 2 | 94.98 | 92.91 | 722.53 |
| MOI30CLC9 | 2 | 20.8 | 22.2 | 60.7 |
| MOI30CLE1 | 2 | 0.26 | 0.19 | 12.24 |
| MOI50CLB9 | 2 | 0.95 | 0.84 | 5.89 |
| MOI50CLE6 | 2 | 100.69 | 98.83 | 28.39 |
| MOI50CLH6 | 2 | 179.64 | 183.06 | 390.82 |
| Average MSE |  | 49.82 | 50.15 | 138.69 |

The results, summarized in Table 5 and Fig 5C, show that also in this scenario the UMI-based method outperformed the MLE of fragment length upon quantifying serial clonal dilutions when measuring distance from the expected value (average MSE 24.2 vs 123.53 respectively). We specifically observed that the relative quantification of the 75% and 50% dilutions datapoints substantially improved using UMI diversity as compared to what previously measured through the more widely used MLE of fragment length. We again noticed that in this experimental setting, the sequencing reads counts showed a quantification accuracy similar to the UMI-based method.

**Table 5:** Expected and observed contribution of the CL6 insertion site upon serial dilution into a bulk HL60 transduced polyclonal population, as measured by IS-Seq using different quantification methods. In the second last row is reported the Mean square error (MSE) between the results of each pipeline iteration and the expected relative abundance of the serial dilutions of CL6. The average MSE for all the results of each pipeline iteration is reported in the last row of the table. (MLE = Maximum Likelihood Estimation, UMI = Unique Molecular Identifiers)

|  | IS-Seq UMI |  |  | IS-Seq reads |  |  | IS-Seq Fragment_MLE |  |  |
| --- | --- | --- | --- | --- | --- | --- | --- | --- | --- |
| Expected (%) | 250 (n=1808) | 500 (n=3513) | 250rep (n=2712) | 250 (n=1808) | 500 (n=3513) | 250rep (n=2712) | 250 (n=1793) | 500 (n=3463) | 250rep (n=2792) |
| 0 | 0.007 | 0.000 | 0.035 | 0.006 | 0.000 | 0.033 | 0.102 | 0.000 | 0.318 |
| 1 | 0.143 | 0.433 | 0.749 | 0.150 | 0.424 | 0.748 | 0.455 | 0.543 | 1.807 |
| 10 | 5.166 | 7.127 | 7.481 | 5.263 | 7.269 | 7.638 | 9.916 | 8.108 | 10.931 |
| 50 | 42.208 | 39.109 | 52.226 | 42.505 | 39.795 | 52.902 | 39.640 | 32.362 | 43.331 |
| 75 | 64.728 | 64.761 | 73.874 | 64.962 | 64.929 | 73.954 | 55.847 | 46.021 | 57.026 |
| 100 | 99.829 | 99.942 | 99.257 | 99.835 | 99.945 | 99.326 | 97.085 | 99.099 | 85.303 |
| MSE | 31.727 | 38.672 | 2.197 | 30.021 | 35.56 | 2.602 | 80.496 | 193 | 97.53 |
| Average MSE | 24.2 |  |  | 22.73 |  |  | 123.66 |  |  |

Because the IS-Seq pipeline uses IS abundance for certain data filtering steps including collision detection, some IS with low abundance might be filtered out in one method but not in the other, hence the total number of integrations obtained using different abundance estimation methods might vary slightly. As an additional control, to account for such differences we merged the datasets obtained using each method for each sequencing run and selected only the identical insertions across the three methods, so to compare the performance of the 3 quantification systems on equal numbers of integration sites. The results shown in Supplementary Table 6,7 confirm our previous findings and show that the UMI-based method has a better accuracy than MLE of fragment length upon quantifying serial clonal dilutions.

### Assessing UMI distribution using experimental and simulated datasets

The results above described suggest that the UMI-based method might be superior to the MLE of fragment length for IS abundance estimation. However, questions remain about whether the UMI distribution as measured upon sequencing in our controlled experiments, was biased towards certain UMI and if this reflected an unbalanced starting library or was instead most likely due to the expected effects of the uneven PCR amplification of vector-genome junctions. To address this point, we first tested the UMI distribution of each sequencing run against the uniform distribution. In order to exclude the effects of the different filtering steps of the IS-Seq pipeline, we artificially extracted the UMIs from the raw R2 FASTQ files using the GTAAGG anchor sequence as reference, therefore measuring the initial UMI content. In Supplementary Table 8, we summarized the number of UMIs obtained in the three sequencing runs described in this study and calculated their relative entropy as described in the Material and Methods section. Notably, despite different sequencing depths, we observed that the UMI diversity was very similar across multiple sequencing runs ranging from 21.788 to 22.263 entropy values.

When visually inspecting the frequency of the most abundant UMIs in each run, we could clearly observe that the distribution of UMIs was already not uniform ahead of all filtering steps of the IS-Seq pipeline (Supplementary Fig.5, left panels). To formally prove this point, we generated for each dataset a uniform distribution by Bootstrapping using the unique UMI extracted from the R2 reads and setting their total number as the sample size base (see Material and Methods for more details). We then calculated the average frequency of all unique UMIs, the entropy of each bootstrapped sample dataset and applied the Kolmogorov-Smirnov(K-S) to compare the observed UMI distribution with the uniform distribution (Supplementary Fig.5, center panels). The results shown in Supplementary Fig.5 (right panels) formally demonstrate that the distribution of UMIs is not uniform. To assess whether this uneven UMI distribution could be due to a preferential and consistent selection of certain UMIs as a result of unbalances in the linker cassette library, we measured the level of sharing of identical UMIs across the 3 sequencing runs. Firstly, we could observe that only 10% of total detected UMI were detected in all runs (Supplementary Fig.6). We then used the recapture of identical UMIs over these 3 sequencing experiments, which were run sequentially at different times, to estimate the total UMI population available for our experiments. By this calculation we could estimate that we had available a total of 14,816,860 different UMIs which correspond to approximately 88% of the maximum theoretical number of different UMI combinations (16,777,216, calculated as 4^^12[combined length of the two random 6nt sequences flanking the 6nt anchor sequence]^) (Supplementary Fig.7). This finding is highly supportive of a very diverse repertoire of UMIs available in our library. To estimate whether the abundance of each UMI in our library could have influenced the abundance of UMI observed in our datasets, we restricted the analyses of shared UMI to the top100 most abundant UMI detected in each sequencing run. The results shown in Supplementary Fig.8 indicates that the level of sharing of the most abundant UMI across the three independent runs is very low (only 2 UMIs are shared among the top 100 most abundant UMIs across the 2 of 3 datasets, but 0 UMIs are shared across 3 data sets), supporting again the notion that we started from a very diverse distribution of UMIs in the original library.

Lastly, we set to model the “real-world” data on UMI diversity described above, with *in silico* simulations to better underscore the effect of different PCR duplicate levels and UMI entropies on IS abundance quantification. Using the simulated data set with 100 IS, we generated different data sets with one ground-truth IS serially diluted in other IS at 100%, 99%, 75%, 50%, 10%, 1%, 0% relative abundance, and tested different levels of artificially created PCR duplicates and entropy levels of UMIs (Supplementary Table 9). As expected, decreasing UMI diversity creates deviations from the true value while decreasing PCR duplicates improved the performance of read-based quantifications. Notably, when we replicated the scenario of our “real-world” library, setting UMI entropy at 14.65927 (resulting from 88% of the UMI combinations available in the simulated library), and modulating the levels of PCR duplicates to up to 200 max reads per IS, we could show that UMI is indeed more likely to outperform not only PCR reads, as expected, but also sonication fragment length abundance. Taken together, the results of our *in vitro* and *in silico* analyses suggest that the library generated using the proposed UMI configuration carries enough diversity to allow for an accurate evaluation of IS abundance in oligo- and poly-clonal experimental settings.

## Discussion

We here described IS-Seq, a new computational pipeline for the identification and quantification of viral integration sites applicable to the study of preclinical and clinical samples. Our pipeline is designed to be used on the Illumina sequencing output from amplicon libraries generated with a specific set of primers and oligos described in the Material and Methods section. However, we showed that it can be adapted to analyze data generated using the INSPIIRED protocol as well.

To test the performance of our pipeline, we designed a set of controlled experiments using single cell clones with known copy numbers and serial dilutions of an individual clone in a polyclonal population. The IS-Seq pipeline also performed efficiently on different in silico simulated scenarios for what concerns precision, recall sensitivity and specificity. However, we believe that the analysis of *in vitro* generated datasets is the most appropriate when it comes to establishing the efficiency and accuracy of a pipeline for IS retrieval and quantification due to the following reasons: 1) differently from the use in silico generated datasets, such method allows testing the efficiency of the computational pipeline in combination with the wet-lab procedure (e.g. efficiency of gDNA fragmentation, binding of the oligos, sequencing library preparation etc.); 2) differently from the use of plasmids designed to reproduce specific vector-genome junctions, it allows establishing the detection and quantification potential of a given pipeline when IS are in their natural genetic and epigenetic milieu (e.g. better for testing the reach of sonication-based gDNA fragmentation or oligos specificity); 3) as compared to other methods, it better reproduces a ”real world” scenario for clonal quantification in that for each cell used as input, the ratio between a given vector-genome junction and the rest of the cellular gDNA is identical.

Throughout this manuscript, we have compared the performance of IS-Seq against INSPIIRED in terms of detection of insertion sites and accurate quantification of their relative abundance, the latter pipeline being the most widely utilized and the most recently described. To this aim we had to adapt either the INSPIIRED or the IS-Seq pipeline to run on each other’s sequencing datasets. We recognize that this reduced our ability to ultimately demonstrate the superiority of one pipeline versus the other, and it should be noted that this was never the scope of such comparison, which is rather intended to show that both pipelines can achieve similar results. Still, in the process, we could highlight certain technical limitations of INSPIIRED when applied to our experimental contexts. For example, we observed that the default configuration of sequence similarity within the BLAT algorithm used by INSPIIRED could be too stringent when applied to certain scenarios such as the one aimed at identifying insertions in immortalized single cell clones. It is possible that either the selective pressure of the single cell expansion or the nature of the genomic content of the HL60 cell line used in these experiments generates a certain degree of sequence divergency with respect to the reference human genome used for mapping, therefore making the 95% sequence similarity threshold too stringent to capture the CL6 insertion. Reducing this threshold alleviated the problem but, concomitantly, increased the amount of background “noise” insertions and substantially expanded the computational time of INSPIIRED (from 2-3 hours with the 95% threshold to 16 hours with the 0% threshold). When applied to these scenarios, the BWA algorithm embedded in the IS-Seq pipeline seems to provide instead more flexibility for IS identification allowing relaxing detection parameters while maintaining the same run time (4 hours on average). While the IS-Seq pipeline retains information relative to sequencing reads mapping in non-unique genomic locations (e.g. repetitive elements), it does not embed them in the final tabular results. We are aware of approaches aimed at addressing these occurrences, for example trying to use sequence similarities among amplicons mapping in repetitive elements to consolidate into individual entities in the attempt to retain more information for clonal tracking purposes (Calabria et al., 2020). We reasoned that, if insertions into repetitive elements constitute an important fraction of the insertion sites pool of a given polyclonal population, missing such events should impact our ability to appropriately quantify serial dilutions of a clone with a known insertion site into a polyclonal cell pool. Specifically, this would result in an over-estimation of the relative abundance of the insertion site of the diluted clone because we would be missing in the denominator several integrations in repetitive elements belonging to our polyclonal population used as dilutant. On the contrary, our results from serial dilution experiments showed that we often tend to underestimate the contribution of our clone of interest with respect to the expected value. In addition, none of the clones with known copy number used in our first test contained insertions in repetitive elements that went missing upon the IS-Seq computational analysis. Indeed, we showed that when we appended reads belonging to putative IS collected using the “multihits” procedure of INSPIIRED to the reads mapping in unique genomic loci, the background noise reached levels that detached the results from the ground truth. These observations lead us to the consideration that, whenever the insertion site retrieval is performed in controlled experimental settings (such as when we can appropriately predict the expected abundance of a given clone over a polyclonal population), the impact of such occurrences seems negligible, therefore we advise extreme caution when attempting to estimate the true contributions of insertions in repetitive elements in samples with unknown clonal composition (such as the ones obtained from patients undergoing gene therapy).

The methods for quantifying the abundance of integration sites have also been objects of debate. Because it is well known that sequencing reads counts could be affected by an uneven exponential amplification of vector-genome junctions during the wet-lab protocol used for insertion sites retrieval, most, if not all, current techniques for insertion sites quantification, now make use of fragment length diversity upon sonication of gDNA. We have previously discussed the theoretical relevance and limitations of such an approach (Biasco et al., 2018) and we could put it to test here against the use of UMIs embedded in the linker cassette. In all our experimental tests, the UMIs largely outperformed the MLE of fragment length, more accurately predicting the contribution of individual clones upon serial dilutions. As predicted in our recent publication (Biasco, 2017), when the starting material ranges from 250ng to 500ng (corresponding to approximately 38,700 and 77,400 human diploid genomes respectively), the available diversity of fragment lengths, even using MLE calculations, is not sufficient to cover for the size of the cell pool composing a clone represented in large numbers over a polyclonal population. Proving this point, the MLE of fragment length largely underestimated the true clonal abundance of the 50% and 75% data points of our dilution experiments, while they became progressively more accurate at bigger clonal dilutions 10-1%. Considering that a 20-30% clonal contribution is normally used for defining a clonal expansion event of clinical relevance, it is important to make use of a method for IS abundance quantification that is as accurate as possible in tracking highly expanded clones. In our controlled experimental settings, the UMI-based quantification approach embedded in the IS-Seq pipeline proved better than the fragment length, for estimating clonal contributions at different dilution levels including at the 50% and 75% single clone contents. In this regard, the analysis of UMI distribution in our datasets strongly suggest that our oligos configuration combined with the IS-Seq sequence processing provides enough UMI diversity and UMI detection accuracy to cover a wide range of clonal contributions, even when applied to the analysis of gDNA samples of the size of the average clinical material (250ng-500ng).

Despite being an efficient tool for IS retrieval and quantification, the IS-Seq pipeline still has some inherent limitations. The current version of IS-seq is implemented in python and R, therefore it requires some proficiency in these languages for the initial setup of the pipeline and for running IS-Seq. Even though we added a “checkpoints” feature to streamline re-runs of the pipeline, we envisage that future versions of this pipeline would benefit from a user-friendly flowchart for keeping track of the entire process, for example, implementing workflow management systems such as Snakemake(Mölder et al., 2021) or Nextflow(DI Tommaso et al., 2017). Moreover, because of the filtering of collisions based on IS abundance, the output of IS-Seq is dynamic in nature and the absolute number of IS per sample can change slightly as new related datasets are added to the analysis, for example, when adding new timepoints from a patient follow up. Should one need to retrospectively report the results of an older ISA data freeze, they must make sure they have archived separately the collision table output of each previous pipeline run. Lastly, because the scope of the manuscript is to evaluate the core function of our pipeline for the detection and quantification of insertion sites, we have not addressed here the implementation of a system for generating reports and downstream analysis of the output of IS-Seq. We specifically designed the format of the output matrixes so that they can be flexibly and easily used as input for further processing. In this regard, our team is using a process based on the *R Markdown* package that generates interactive html documents whose content is fully customizable with tables and figures and whose code and use is described in our GitHub repository (see Material and Methods).

In summary, we here described IS-Seq, a bioinformatics tool for IS identification and quantification based on the use of different abundance estimation methods embedded in a computationally efficient and automated pipeline. We could show that IS-Seq performs as well as INSPIIRED, the current benchmark pipeline for IS identification, therefore expanding the toolbox available to researchers for the clonal tracking of vector-marked cells upon preclinical and clinical gene therapy.

## Materials and Methods

### Generation of single cell clones

To generate single cell clones for our experimental datasets, we made use of the HL60 cell line (ATCC, CCL-240^TM^). On Day 0, HL60 cells were seeded at 1,000,000 cells/mL with IMDM in the presence of 20% FBS and 1% Pen/Strep in 6-well plates. HL60 WT cells were then transduced with a lentiviral vector encoding for the GFP marker gene at the following Multiplicity Of Infection (MOI) of 5,10,20,30 and 50. After 1 day the medium of the transduced HL60 cells was replaced with fresh IMDM and the cells were then kept in culture for 14 days. On Day 14, half of the transduced HL60 cells were pelleted and frozen to be analyzed by ddPCR for their average VCN. To generate single cell clones, we sorted the HL60 transduced cells using the SONY MA900 sorter. Briefly, cells were stained with DAPI to discriminate between living and dead cells. A total of 192 cells with high GFP MFI were sorted individually into two 96-well plates in 100 μL IMDM complete media. Cells were then cultured in 96-well plates and further expanded in 24-well plates for 27 days, after which time the resulting single cell clones were pelleted for molecular analyses. For the clonal dilution experiment, the gDNA from clone CL6 with average VCN=1 was diluted into gDNA from the transduced polyclonal HL60 bulk population at different relative contributions (100%, 75%, 50%, 10%, 1% and 0%) for a total amount of starting material of either 250ng or 500ng, which was then used for insertion sites retrieval, sequencing and downstream analysis.

### VCN evaluation by ddPCR

Genomic DNA (gDNA) isolation from single cell clones was performed using a QIAamp 96 DNA QIAcube HT Kit (Qiagen). DNA concentrations were measured by Qubit™ 4 Fluorometer (Life technologies) using Qubit® dsDNA BR (Broad-Range) Assay Kits (Life Technologies). The VCN assay was performed by ddPCR using 1) Vector-specific oligos designed on the Woodchuck Hepatitis Virus Posttranscriptional Regulatory Element (WPRE): [Forward: TTCTGGGACTTTCGCTTTCC, Reverse: CCGACAACACCACGGAATTA, Probe: 5’ FAM/ATCGCCACGGCAGAACTCATCG/3IABkFQ] and 2) oligos designed on the Glycosyltransferase-like domain-containing protein 1 (GTDC1) as human gene control [Forward: GAAGTTCAGGTTAATTAGCTGCTG, Reverse: GGCACCTTAACATTTGGTTCTG, Probe: 5’ HEX/ACGAACTTCTTGGAGTTGTTTGCT/3IABkFQ]. The ddPCR mix was prepared using 10 µL of ddPCR Supermix for Probes No dUTP (BioRad), reverse and forward primers and probes at final concentrations of 100 uM, 30 ng of gDNA and nuclease-free water in a total volume of 20 µL. The mixes were then loaded into the DG8™ Cartridge, and droplets were generated automatically with the QX200™ Droplet Generator. The emulsion was transferred to a PCR plate and cycled using the following thermal cycler conditions: Hold: 95°C for 10 min, 40 cycles at 95°C for 30 s, 60 °C for 1 min, then enzyme deactivation 98 °C for 10 minutes and final hold at 4 °C. Data acquisition and analysis were performed using the QX200™ Droplet Reader and QuantaSoft™ Software (Biorad).

### Generation of the simulated data set based on the IS-Seq design

We developed a procedure to generate an *in silico* simulated dataset based on the FASTQ output of the IS-Seq wet-lab analytical pipeline. The generation of the simulated datasets was based on the following steps: 1) we first set an integration site position to define the ground truth (to this goal we used the chr19, 49461738, - strand position of the CL6 IS); 2) next we extracted a 3kb DNA Fragment (chr19: 49458738 _49461738) from the positive strand of the host reference genome (hg38ChrOnly.fa). Since this IS is in the negative strand, we took the reverse-complement of this 3kb as the 3000_fragment input FASTA for the simulation; 3) to run the simulation we used the ART software(Huang et al., 2012) with the following settings: *art_illumina -ss MSv3 -p -I 3000_fragment.fa -l 250 -c 17274461 -m 1000 -s 300 -d “simulate” -o simulationUp_3000_Frag*; 4) We next aligned the simulationUp_3000_Frag1.fq and simulationUp_3000_Frag2.fq output files to the human genome (hg38ChrOnly.fa) using the following command: *bwa mem2 -t 8 path/to/hg38ChrOnly.fa path/to/ simulationUp_3000_49461738_Frag1.fq path/to/ simulationUp_3000_49461738_Frag2.fq > path/to/ UpFrag49461738R1_R2_Barcode_FB-P5-Rd1-LTR.9_FB-P7-Rd2-LC.9_aligned_mem.sam*; 5) To follow, we run sam2filterNo.R on path/to/ UpFrag49461738R1_R2_Barcode_FB-P5-Rd1-LTR.9_FB-P7-Rd2-LC.9_aligned_mem.sam to get POOL-ISA-AVRO-6-Preclin_FB-P5-Rd1-LTR.9_FB-P7-Rd2-LC.9_final_parse_filterNo.txt file; 6) Then, from the * final_parse_filterNo.txt file generated in step5, we obtained the read IDs that aligned to the chr19, 49461738, -strand position; 7) Using the read IDs identified in step6, we selected the reads that match to these IDs from the simulationUp_3000_Frag1.fq and simulationUp_3000_Frag2.fq files; 8) Based on the R1 reads selected in step7, we created new R1 reads appending the IS-Seq Illumina adapter sequence, sample barcode, LTR to the 5’ end of R1 in the following order: Illumina adapter sequence(12nt) + sample barcode(8nt) + LTR(32nt) + R1 Similarly, based on the R2 reads selected in step7, we created new R2 reads appending the IS-Seq Illumina adapter sequence, sample barcode, LC1, UMI, LC2 to the 5’ end of R2 in the following order: Illumina adapter sequence(12nt) + sample barcode(8nt) + LC1(29nt) + UMI(18nt) + LC2(20nt) + R2. Note: The UMI sequence includes 6nt random sequence + 6nt anchor sequence + another 6nt random sequence.; 9) Lastly, we updated the read ID of these new R1 and R2 reads as “Mchr19-simulate34517884 1”, and saved these new R1 and R2 reads as simulationUp_R1.fq.gz and simulationUp_R2.fq.gz; 10) The *.fq.gz files generated in step9 were then used as input for running IS-Seq. The steps 1-10 are used to obtain the reads in the form of LTR-host genome fragment junction for the 1, 5, 100 or 1000 integration sites we decided to set up as the ground truth. In order to measure the performance of IS-Seq on different simulated scenarios, we further introduced mutations in the first genomic 50nt after the LTR with 10% reads of R1 reads and added 17,274,461 random reads as background noise to evaluate the ability of IS-Seq to discriminate true IS from random sequencing reads. To generate the dilution data set with one IS diluted at 100%, 99%, 75%, 50%, 10%, 1%, 0% relative abundance into other IS, using the simulated data set with 100 IS, we considered chr19_49461738_-as the true IS, and set different levels of PCR duplicates and UMI entropies to obtain different simulation data sets. We then calculated the MSE between the relative abundance of different simulation scenarios vs the ground truth value artificially established. The results of these analyses are described in the results section and shown in Supplementary Table 4, 9 and Supplementary Figure 9.

### Adaptation of the INSPIIRED and IS-Seq pipelines for comparative analyses

For running INSPIIRED on the IS-Seq datasets, we re-implemented 3 INSPIIRED key functions, namely *demultiplex*, *trim_after_demultiplex* and *integration_site_calling*. First, we tested our INSPIIRED implementation on the INSPIIRED data set from the relative GitHub repository. For this test, a user needs 3 input FASTQ files(*I1_001.fastq.gz,* R1_001.fastq.gz, *R2_001.fastq.gz), one sample information file (sampleInfo.tsv), one parameter file (processingParams.tsv) and a vector sequence file (p746vector.fasta). *I1_001.fastq.gz contains the sample barcode index, * R1_001.fastq.gz, *R2_001.fastq.gz include reads needed to be demultiplexed for all samples. *I1_001.fastq.gz is used as an input for demultiplex.R. The I1_001.fastq.gz file is trimmed by trimTailw function in ShortRead R package based on the quality of reads, and further selected to keep reads with 12nt. The selected index reads are decoded using the *decode* function from python, and the reads with the correct code are selected in the processGolayTest.py script. Then, the reads of each sample are demultiplexed to each sample by matching the read name in the index file with the read name of two FASTQ files. Sample information, parameter, vector sequence, and the location of demultiplexed FASTQ files are processed and stored into a completeMetadata file, and this file is also used as an input for the *demultiplex.R* function. After this step, the demultiplexed paired-end reads of each sample are used as input for the *trim_after_demultiplex* function for the steps of filtering R1 and R2 reads, matching R1 and R2 and aligning to host reference genome by blat under the parameter setting in the completeMetadata file. The aligned *R1.psl and *R2.psl files are then used as input for the *integration_site_calling* function to call IS. After finished calling IS, we use the allSites.rds file from the output files to get the fragment length count data. Lastly, these fragment counts are used to estimate fragment MLE using the *sonicLength* R package.

When we run our INSPIIRED implementation on the data set from IS-Seq, we started from the R1 and R2 FASTQ files (R1*fq_trimwithCutAdapt and R2* fq_trimwithCutAdapt FASTQ files) that were already processed by the IS-Seq pipeline (after demultiplexing, trimming and filtering steps). R1*fq_trimwithCutAdapt and R2* fq_trimwithCutAdapt files are selected by matching read name with R1_R2_Barcode*_trimmedID (an output file from the IS-Seq pipeline) to generate R1*fq_trimwithCutAdapt_ReadyToAlignSort and R2*fq_trimwithCutAdapt_ReadyToAlignSort files. These *ReadyToAlignSort files are converted into FASTA files by fastq_to_fasta function from FASTX Toolkit. The converted R1*.fa and R2*.fa are then aligned to the host reference genome using BLAT to generate R1*.fa.psl and R2*.fa.psl files; all these steps are implemented in the *FqToFa.R* script. R1*ReadyToAlignSort and R2*ReadyToAlignSort are matched by read name to generate a keys.rds file by MakeKeys.R Script. The aligned R1*.fa.psl, R2*.fa.psl files and key.rds files are subsequently used as input for the script called *PslToIs_one_replicate_change_sequence_similarity.R* to call the IS. Due to the different oligo design, in the protocol for IS retrieval used for running INSPIIRED, the LTR sequence is on R2, while in the wet-lab protocol at the basis of IS-Seq the LTR is on R1. Therefore, in this step, we had to swap R1 with R2. We used R2*.fa.psl as R1.psl and R1*.fa.psl as R2.psl to call integration sites and abundance quantification with INSPIIRED. Other inputs for *PslToIs_one_replicate_change_sequence_similarity.R* include completeMetadata.RData, the user-defined output directory, which reference genome is to be used, sample index and sequence similarity threshold for filtering reads from the aligned *.psl files (for this manuscript we used 95%, 80% and 0% thresholds). Lastly, fragments MLE are obtained by applying the *sonicLength* R package on the fragment length counts extracted from the allSites.rds output file. Because we discovered that for efficiently detecting the CL6 insertion site INSPIIRED needed to be run at SST=0%, for a fair comparison with our pipeline, and to give INSPIIRED a better chance, we also run IS-Seq with MAPQ=0.

Conversely, to apply IS-Seq to the INSPIIRED dataset, we needed to convert the input file format of the INSPIIRED dataset to a format recognizable by IS-Seq. To this aim, we implemented the following procedures. We first converted the *I1_001.fastq.gz, *R1_001.fastq.gz, *R2_001.fastq.gz files into R1. fastq.gz and R2_001.fastq.gz files. In this step, *I1_001.fastq.gz is selected by matching the read name in the correct decoded FASTA file (correctedI2-1.fasta), and the selected reads are matched with the reads in * R1_001.fastq.gz, *R2_001.fastq.gz files. To follow, we appended each sample barcode of the index file at the beginning of the related R1 and R2 reads. Due to the different oligo design between INSPIIRED and IS-Seq we mentioned above, after appending sample barcode, *R2_001.fastq.gz is outputted as R1.fq.gz, and *R1_001.fastq.gz is outputted as R2.fq.gz. Next, the barcode_LTR.fil and barcode_LC.fil files were generated based on the values of the bcSeq column of the sampleInfo.tsv file from the INSPIIRED Github repository(https://github.com/BushmanLab/intSiteCaller/blob/master/testCases/intSiteValidation/sampleInfo.tsv). Then, we generated the LTR_lentiviral.fa combining the primer and ltrbit columns of sampleInfo.tsv, the LC_completo.fa from the leftmost 19nt sequence of linkerSequence column in sampleInfo.tsv (the 4 replicates of each clone would share the same sequence, but different clones would have different LC_completo.fa sequence), the LC1brcd_ancora.fa (as an empty file, since INSPIIRED has no anchor sequence embedded in the UMI) and the LC1brcd_last.fa from the rightmost 16nt sequence of the linkerSequence column in sampleInfo.tsv (all samples would have the same sequence). Lastly, we generated one association file for each clone and downloaded the hg18 as human genome reference as well as the vector sequence FASTA file and their bwa index files. After all these files had been generated, we could use them as input to run IS-Seq starting from the initial step of the pipeline.

### Extracting UMI and checking UMI distribution using IS-Seq

In the IS-Seq pipeline, we use UMI-tools (Smith et al., 2017) to process UMI tags. The UMI-based abundance estimation is based on these extracted UMI-tags. To check UMI distribution, we develop our own R script to extract the UMIs from R2.fq.gz file using the GTAAGG anchor sequence embedded in each UMI tag. When browsing for the 6nt anchor sequence to identify the UMI region on R2, we can occasionally get secondary hits matching to the human genomic region flanking the linker cassette. Therefore, IS-Seq selects the hit position with the highest match frequency and double checks that it corresponds to the expected position in R2 to extract the UMI sequence. We defined these matched UMI sequences as “right-matched” UMIs and used an entropy measurement to quantify the randomness of the distribution of these UMIs. The entropy is calculated using the following formula:

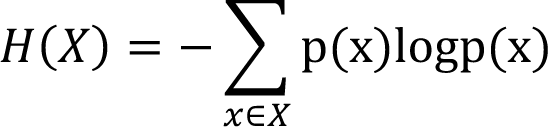

Where X is the right-matched UMI set, x is one UMI in the UMI set and p(x) is the occurrence probability of x in the UMI set.

To generate a uniform UMI distribution for each data set and to compare it with the real UMI distribution, we first extracted the unique UMIs from the total right-matched UMIs. We then performed 100 Bootstrap sampling using the unique right-matched UMI and set the number of right-matched UMI as the base of sample size (note: for the single clone and the 250rep dilution dataset, we used the number of right-matched UMI, but for the dilution dataset 250-500, we had to multiply by 10fold the number of right-matched UMI to be used as sample size given that we would have ended up sampling 5,069,672 UMIs from 4,180,988 UMIs, hence under-representing the UMI population). From the 100 bootstrapped samples, the average occurrence frequency of each unique right matched UMI was calculated and used as uniform distribution for the null hypothesis. The entropy of each bootstrapped sample was also calculated and used as entropy distribution for the null hypothesis. Lastly, we used the Kolmogorov-Smirnov(K-S) test to compare data UMI distribution with the uniform UMI distribution.

To estimate the UMIs population size, we used data derived from the three independent sequencing runs generated for this manuscript chronologically ordered by date of sequencing as 1) dilution dataset 250-500, 2) single cell clone dataset and 3) dilution dataset 250rep. We then applied the capture-recapture method embedded in the *Rcapture* R package (Baillargeon and Rivest, 2007; Ramos et al., 2020), and used the *closedp.mX* function to fit a loglinear model for 3 capture occasions without including any interaction.

### Details, installation and use of the IS-Seq pipeline

A detailed description of the IS-Seq pipeline is available at the following repository https://github.com/aiminy/IS-Seq-python3 and includes a full tutorial on how to install and use this tool as well as all the necessary files and input templates. We created a Docker image based on Python 3.9.12 and R 4.1.2 with all software, tools and R packages required to use IS-Seq to facilitate the setup of this pipeline for non-expert users. We also supplied a tutorial to show how to use IS-Seq on the simulated data set including 1 IS. In this manuscript we used BLAT to align R1 and R2 IS-Seq reads to the host reference genome when adapting for INSPIIRED, using the same BLAT parameters of INSPIIRED for consistency. In the IS-Seq pipeline, we supply an additional option, also based on BLAT, to allow the user to check how many reads align to a given vector sequence. The parameters used for this procedure are based on the standard configuration of BLAT for identifying sequence match on the UCSC genome browser.

## Data availability Statement

Data sets will be available through GEO when this manuscript is accepted for publication.

## Author contributions

AY implemented the pipelines, performed the computations and analysis and drafted the manuscript. CB, AN, LB and ML performed experiments and generated all sequence data sets. LB supervised the project, wrote the manuscript and designed Figures for visualization.

## Acknowledgments

The version of the IS-Seq pipeline presented here is the result of a long effort spanning several years with contributions from multiple researchers. We would like to thank in particular Lorena Leonardelli for the initial design of the modules composing IS-Seq and Danilo Pellin for assembling and optimizing an early version of the automated pipeline. We are also grateful to Andrea Calabria and Raffaele Fronza for the many scientific conversations about integration sites detection systems and analysis. The wet-lab protocol at the basis of IS-Seq is also based on important inputs from several actors including Fabrizio Benedicenti at San Raffaele Telethon Institute for Gene Therapy and Frederic Bushman at UPENN. We also want to acknowledge researchers at University College of London and AVROBIO, Inc. for the support and the many productive scientific discussions. Lastly, this manuscript is dedicated to the memory of Manfred Schmidt, who pioneered insertion sites analysis and who has been a mentor and a friend.

## Supplementary figures

**Supplementary Figure 1:**
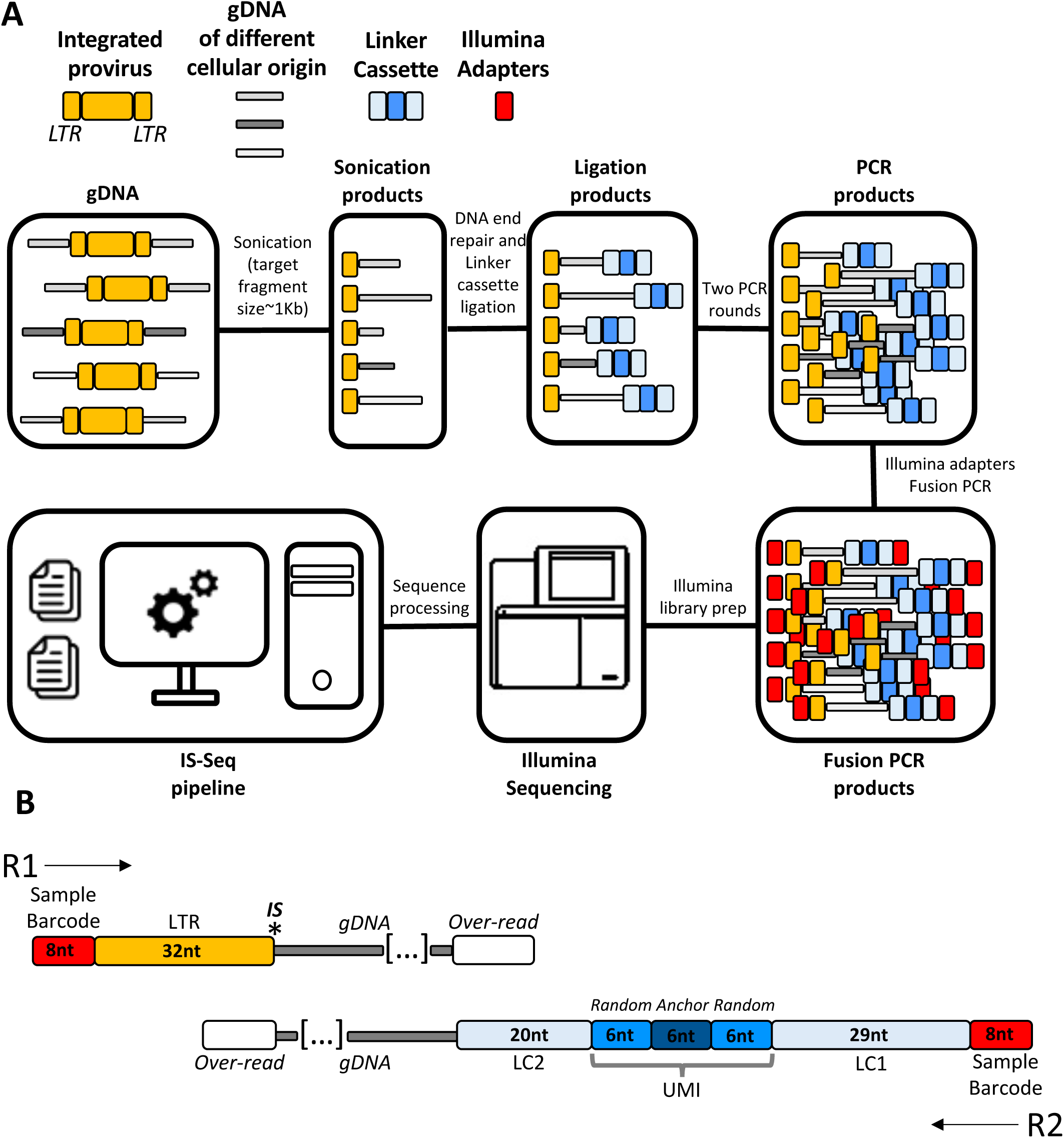
A) Schematic of the procedure used for insertion site retrieval. B) Detailed view of the sequence structure of R1 and R2 upon Illumina sequencing (does not include Illumina adapters, P5 and P7 and the 12nt random sequence used to facilitate cluster recognition). (LTR=Long Terminal Repeat, IS= Insertion site, LC1 = Linker cassette segment 1, LC2 = Linker cassette segment 2, UMI = Unique Molecular Identifier, nt = nucleotides)

**Supplementary Figure 2:**
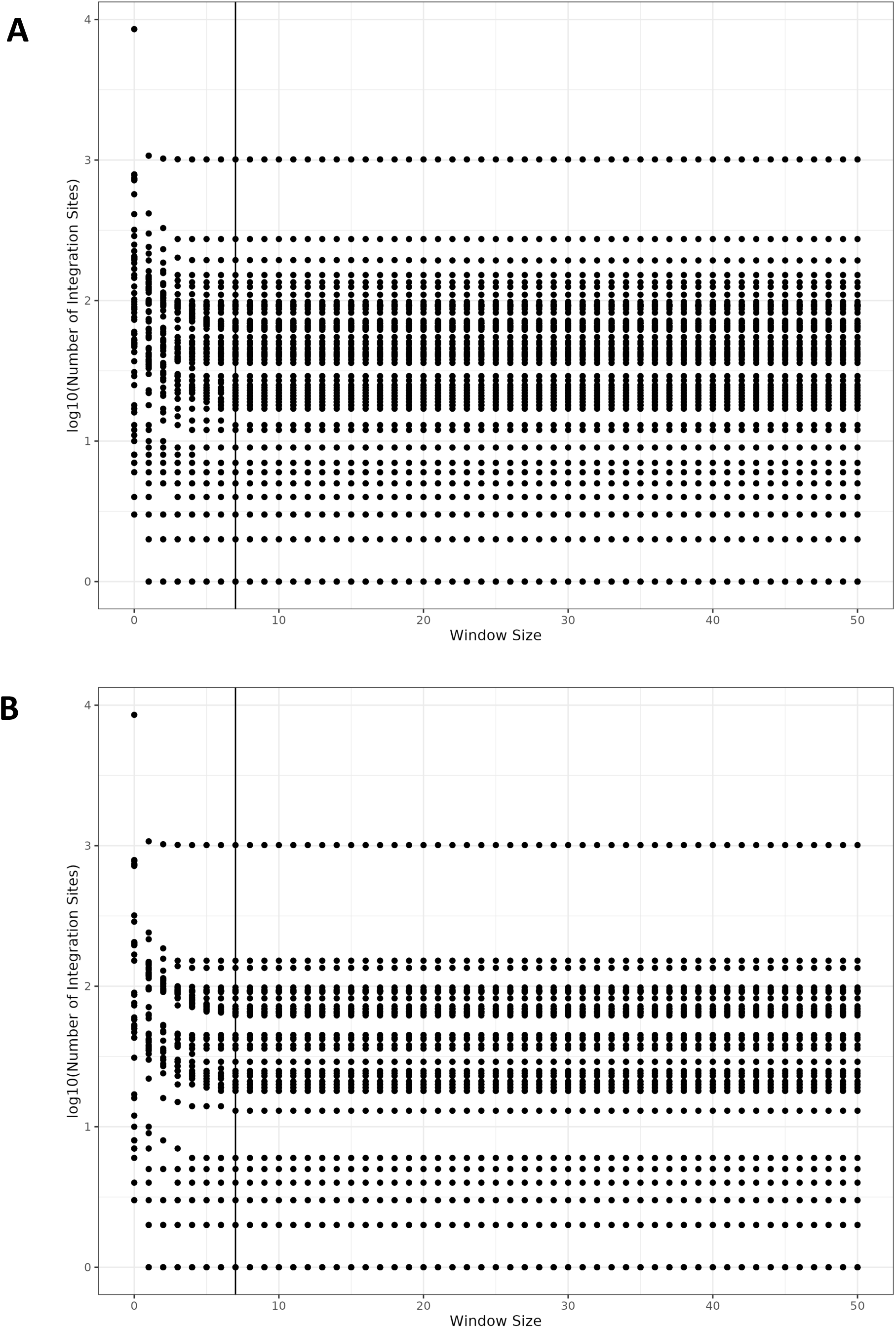
Number of integration sites detected in all samples and datasets analyzed in this manuscript using different windows for IS merging. Datasets reported are a combination of the outputs of filterNo and filter60 (A) or filter60 only (B). Each dot in the plot represent a sample, vertical line indicates the 7bp window.

**Supplementary Figure 3:**
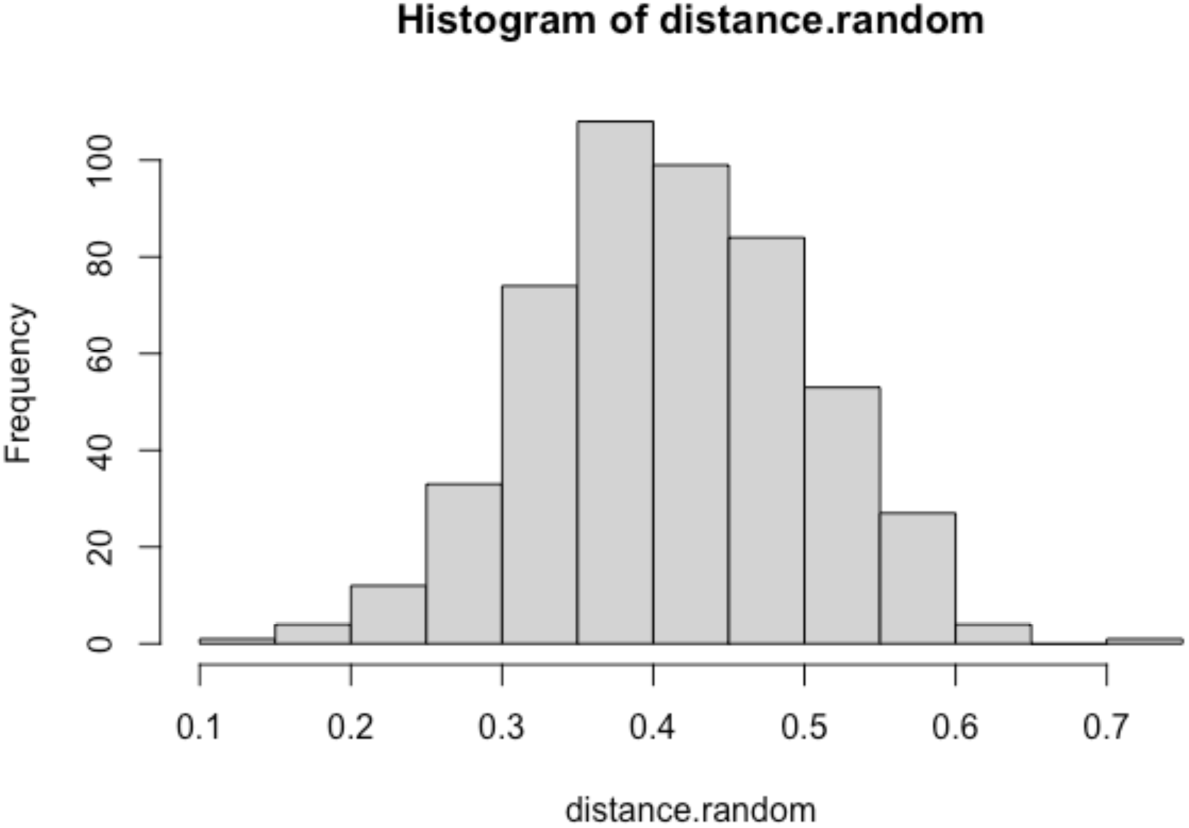
Frequency distribution of distances between INSPIIRED relative abundance readouts and 500 matrixes randomly generated from the IS-Seq relative abundance results.

**Supplementary Figure 4:**
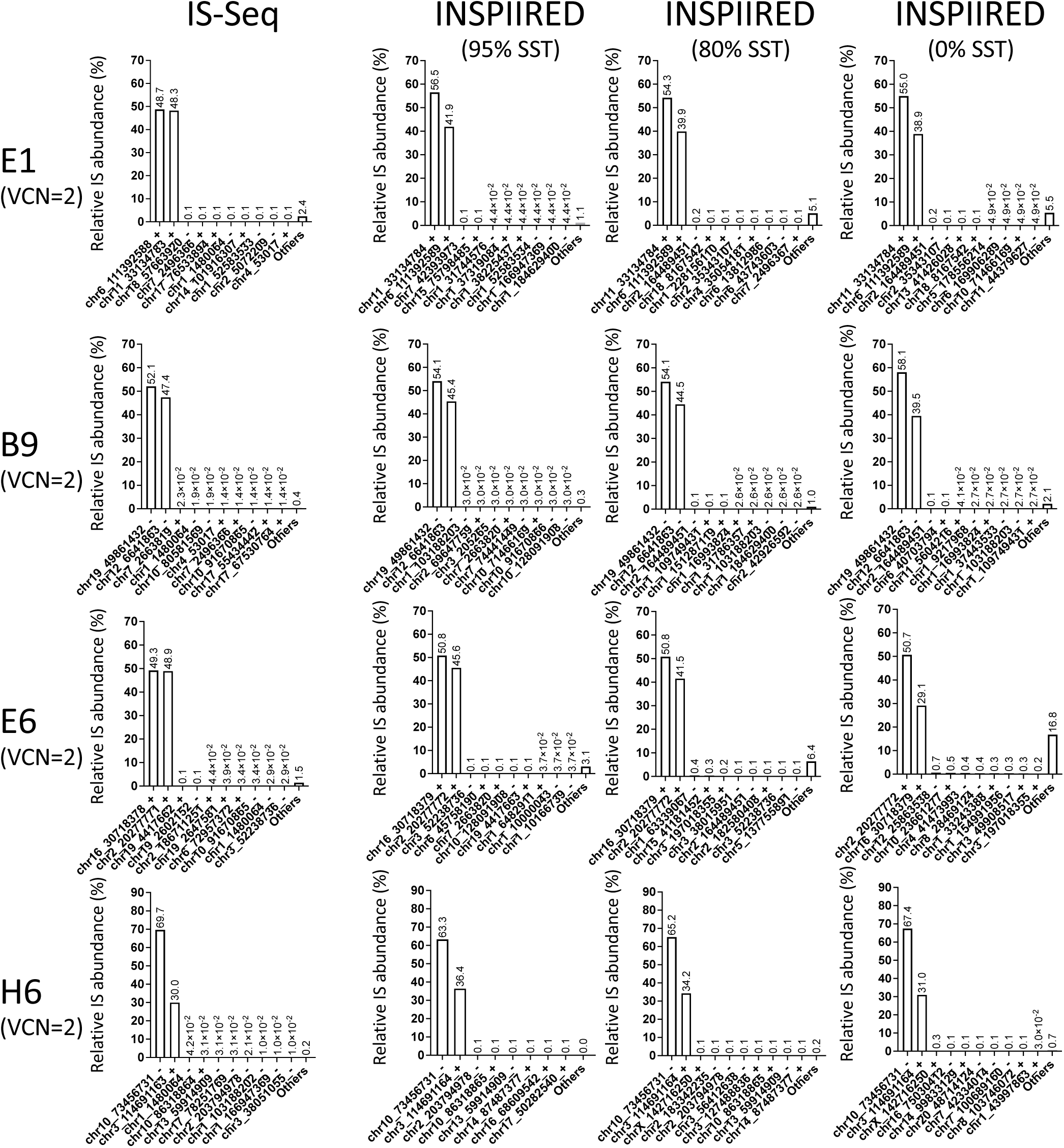
Relative abundance of insertion sites (IS) detected in single cell clones using different pipelines and sequence similarity thresholds (SST) based on the Maximum Likelihood Estimate of fragment lengths. (VCN = vector copy number) (additional results in Fig. 4)

**Supplementary Figure 5:**
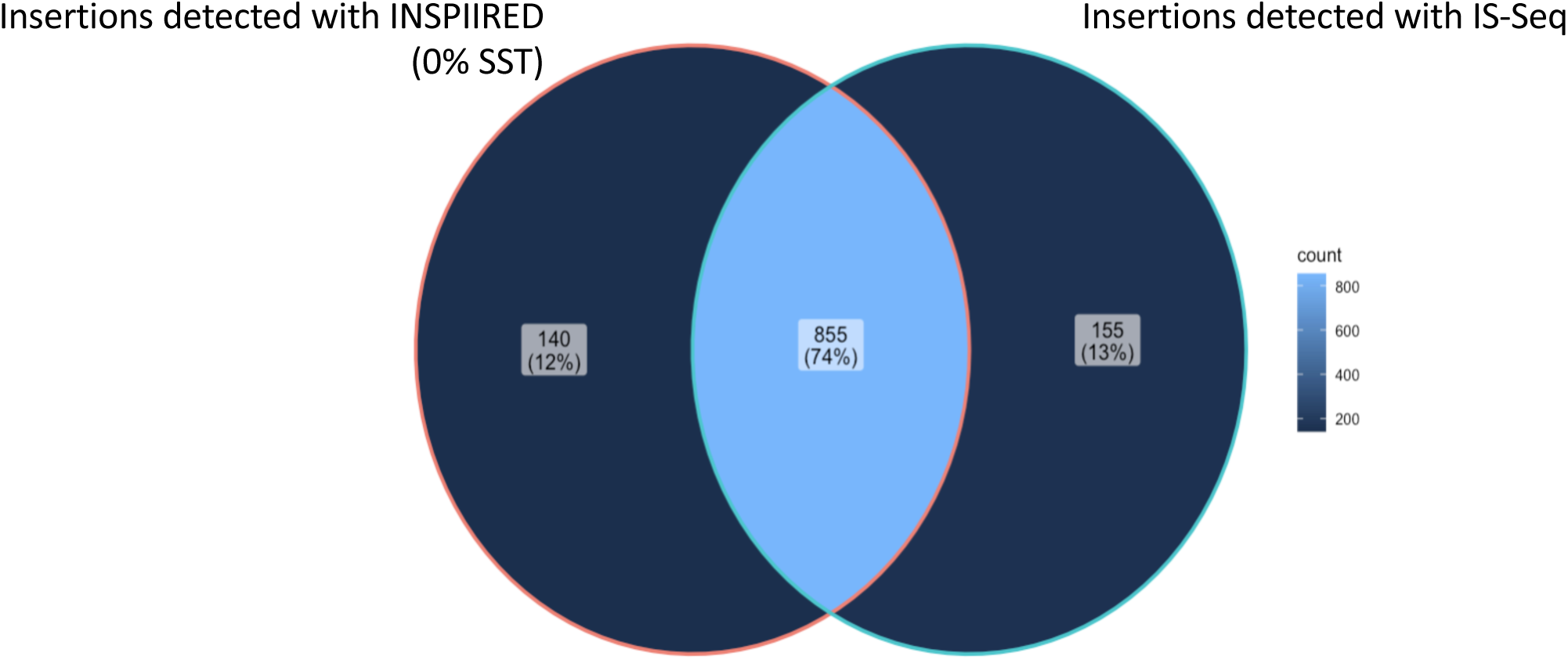
Sharing of identical insertions detected with INSPIIRED and IS-Seq in the HL60 polyclonal bulk population (0% dilution point of the CL6 clone) (SST = sequence similarity threshold)

**Supplementary Figure 6:**
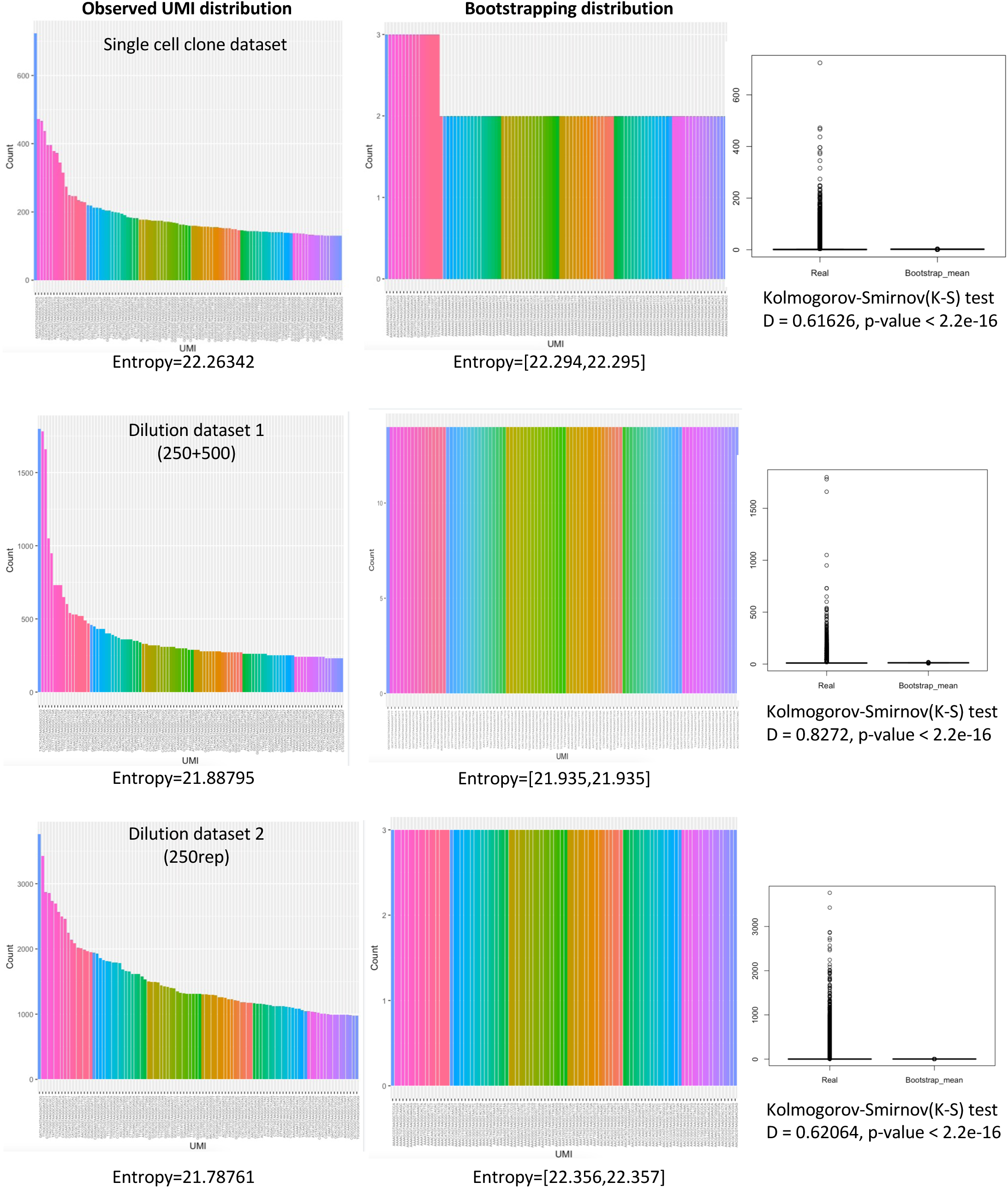
Left panels show the distribution of the top 100 most abundant UMIs reads counts observed in the 3 datasets object of this study. Each bar represents an individual UMI whose sequence is reported in the x-axis. Below each plot is reported the entropy value of the whole UMI population. The center panels show the same analysis performed on 100 a random subsampling (bootstrapping) of UMIs as described in the main text. Range of entropy values is reported below each plot. The right panels show the global distribution of the observed UMIs counts (real) vs the mean of the counts obtained with bootstrapping. Kolmogorov-Smirnov(K-S) test results are shown below each plot (D = distance).

**Supplementary Figure 7:**
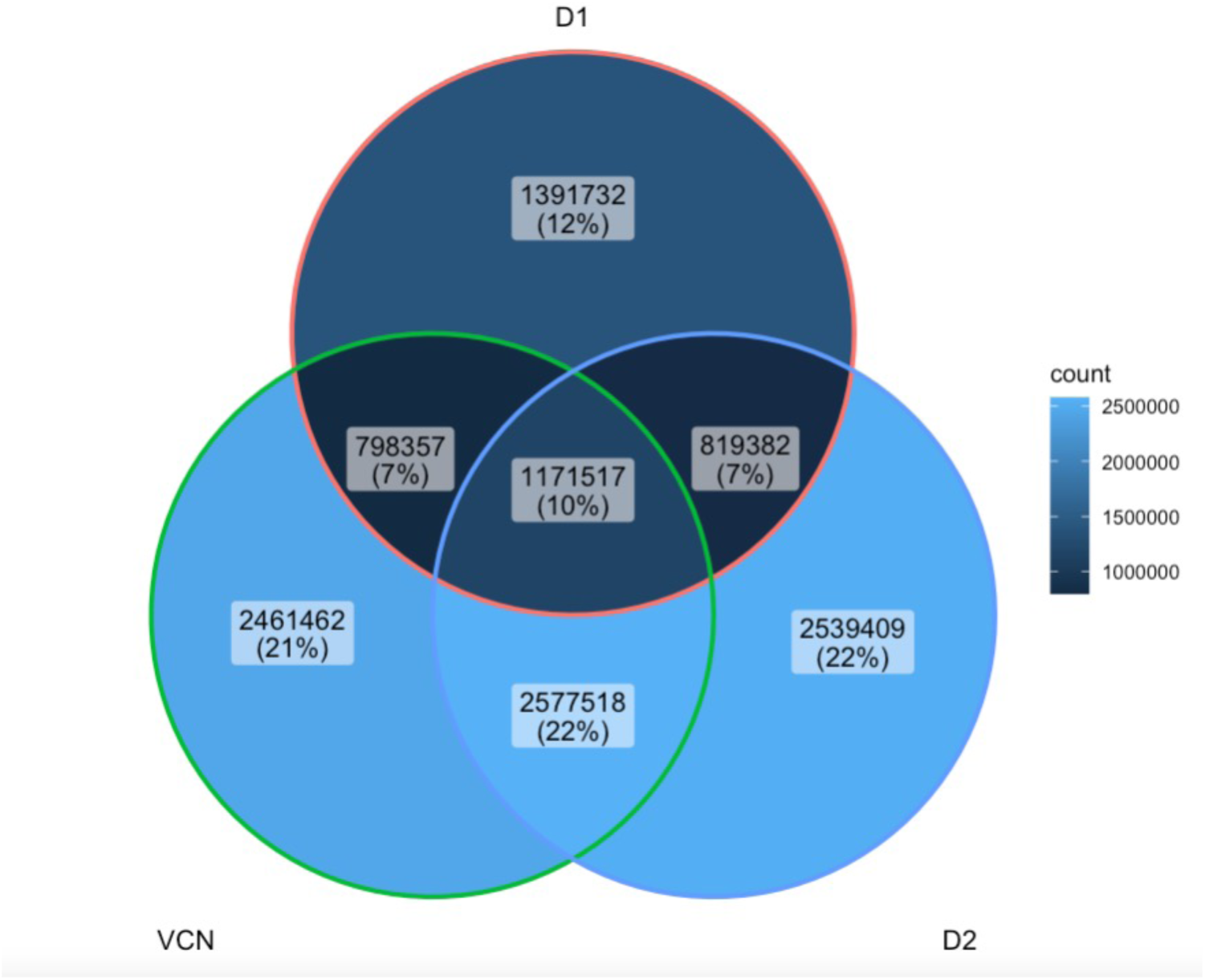
Sharing of identical UMIs among the 3 datasets object of this study.

**Supplementary Figure 8:**
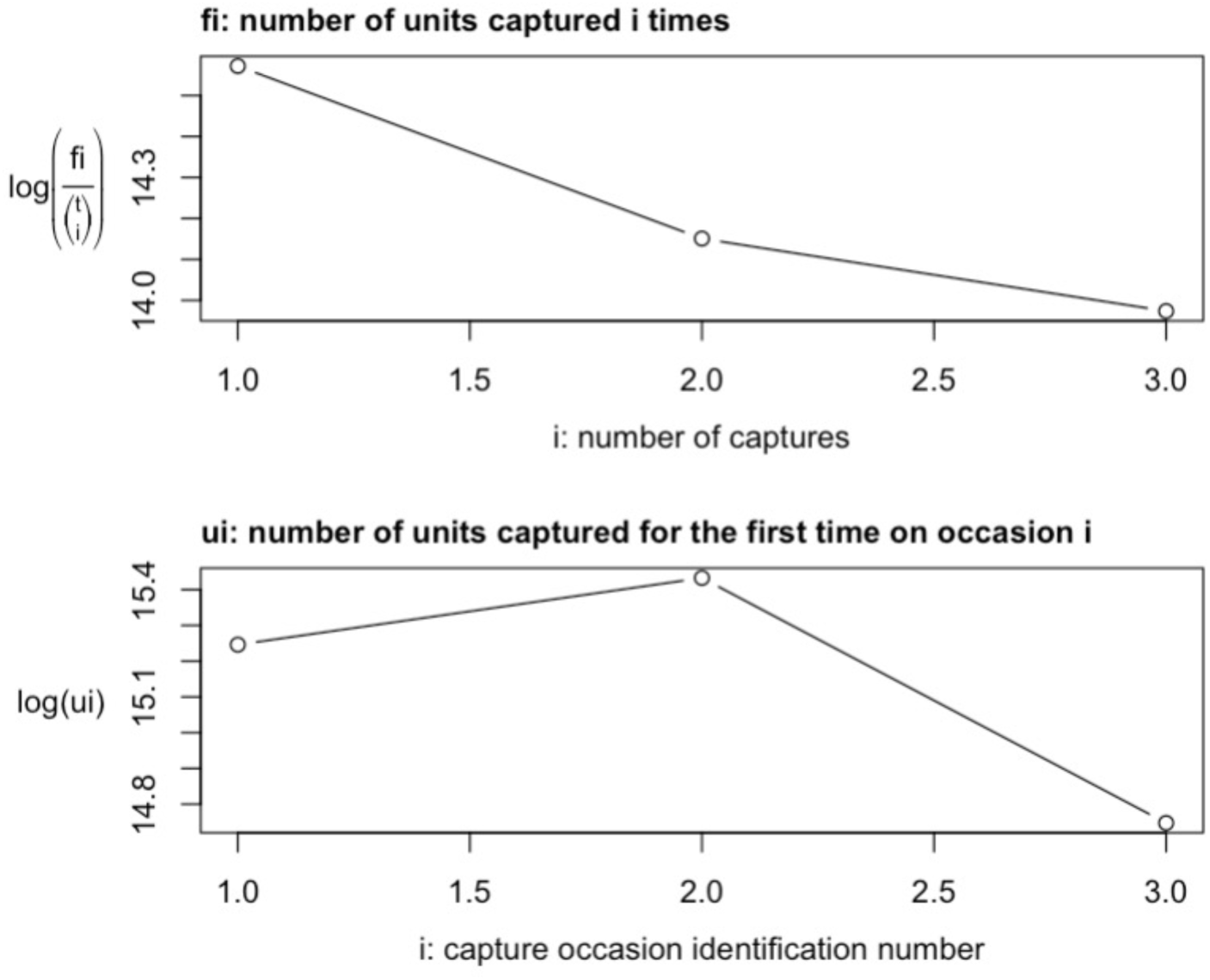
UMIs recapture over 3 independent sequencing runs chronologically ordered based on date of sequencing where 1=dilution dataset 1 (250-500), 2= single cell clones dataset and 3= dilution dataset 2 (250rep). The top panels show the number of units re-captured overtime while the bottom panel the number of new units captured at each sequencing run.

**Supplementary Figure 9:**
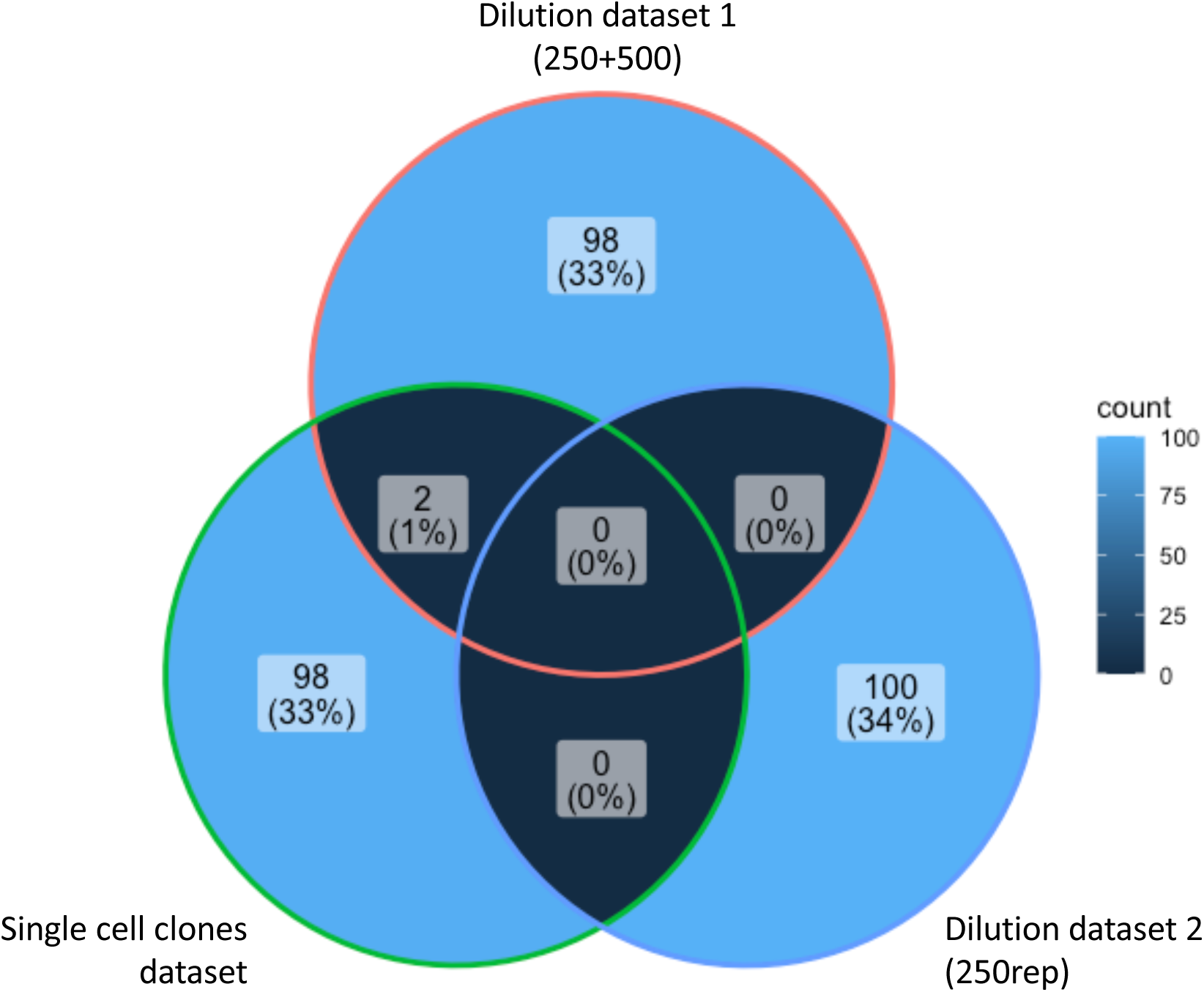
Sharing of identical UMIs among the most abundant top 100 UMIs detected in the 3 datasets object of this study.

**Supplementary Figure 10:**
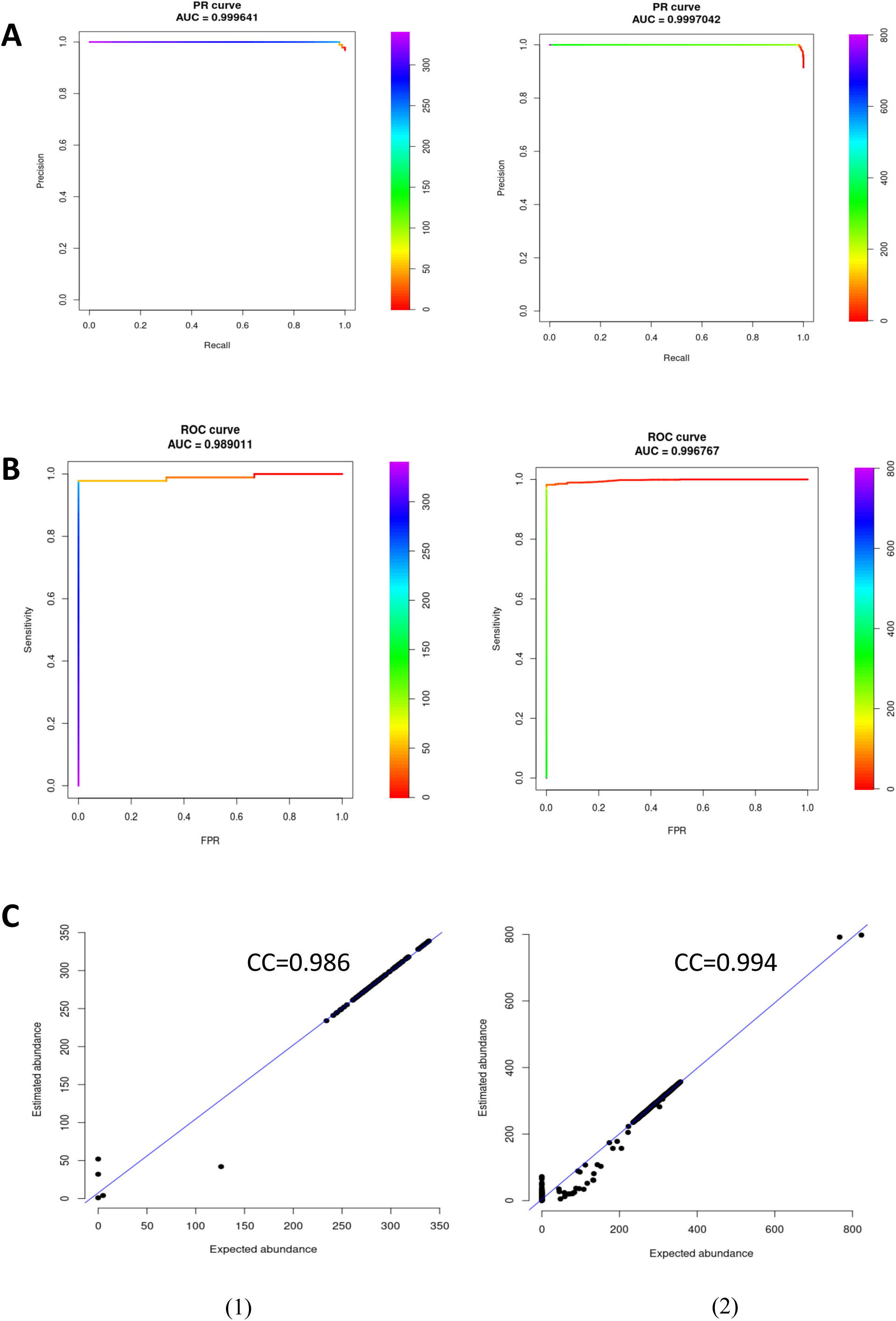
IS-Seq performance metrics based on Precision-Recall curve (A), ROC curve (B) and correlation between expected abundance and estimated abundance (C) on simulated datasets with 100 (left panels) and 1000 (right panels) integration sites (AUC= area under the curve, CC= correlation coefficient).

**Supplementary Figure 11:**
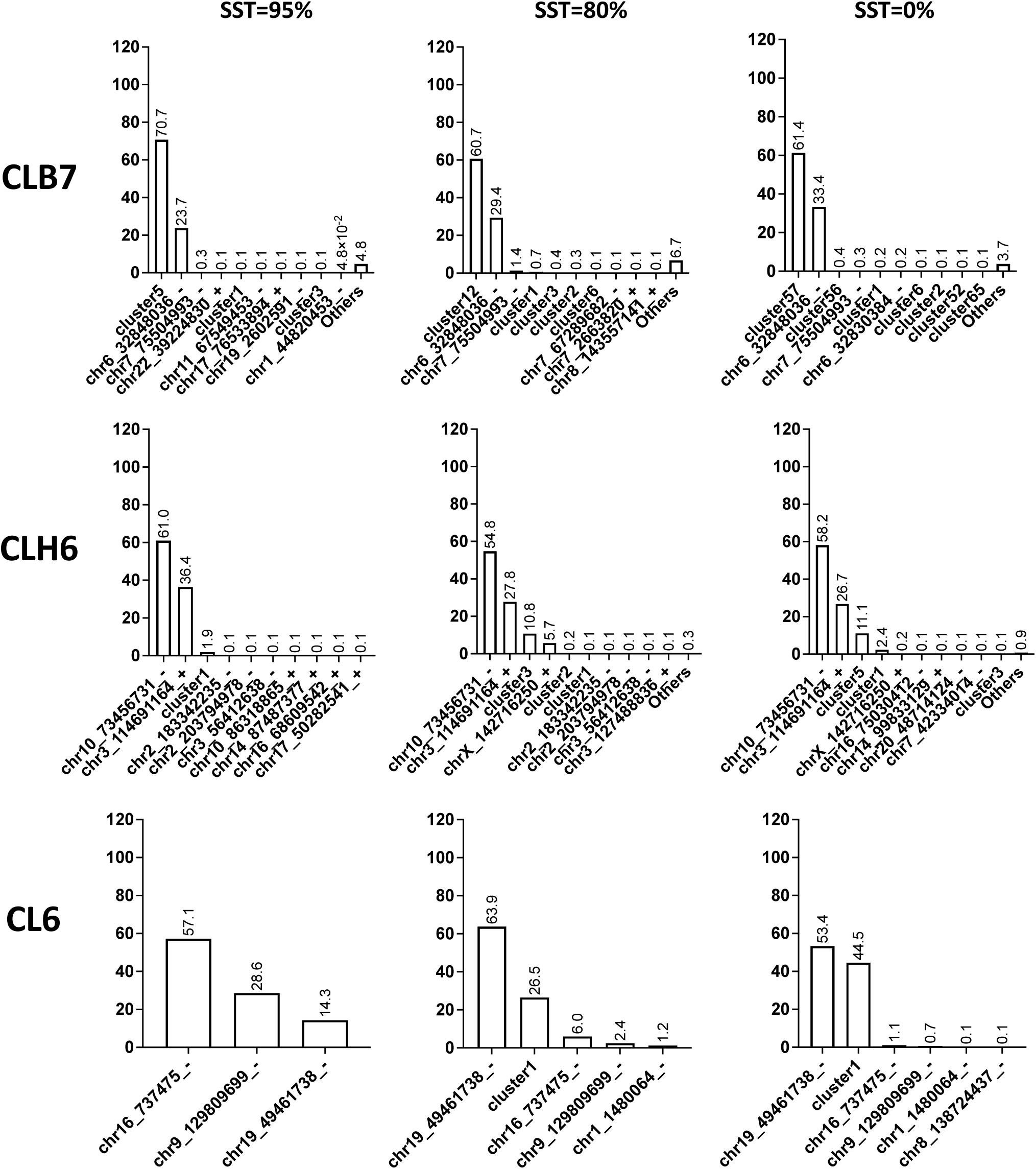
A) Relative abundance of MLE of fragment lenghts of IS collected from CLB7, CLH6 and CL6 using INSPIIRED combining unique hits and sequence clusters from the multihits at different sequence similarity thresholds (SST).

**Supplementary Figure 12:**
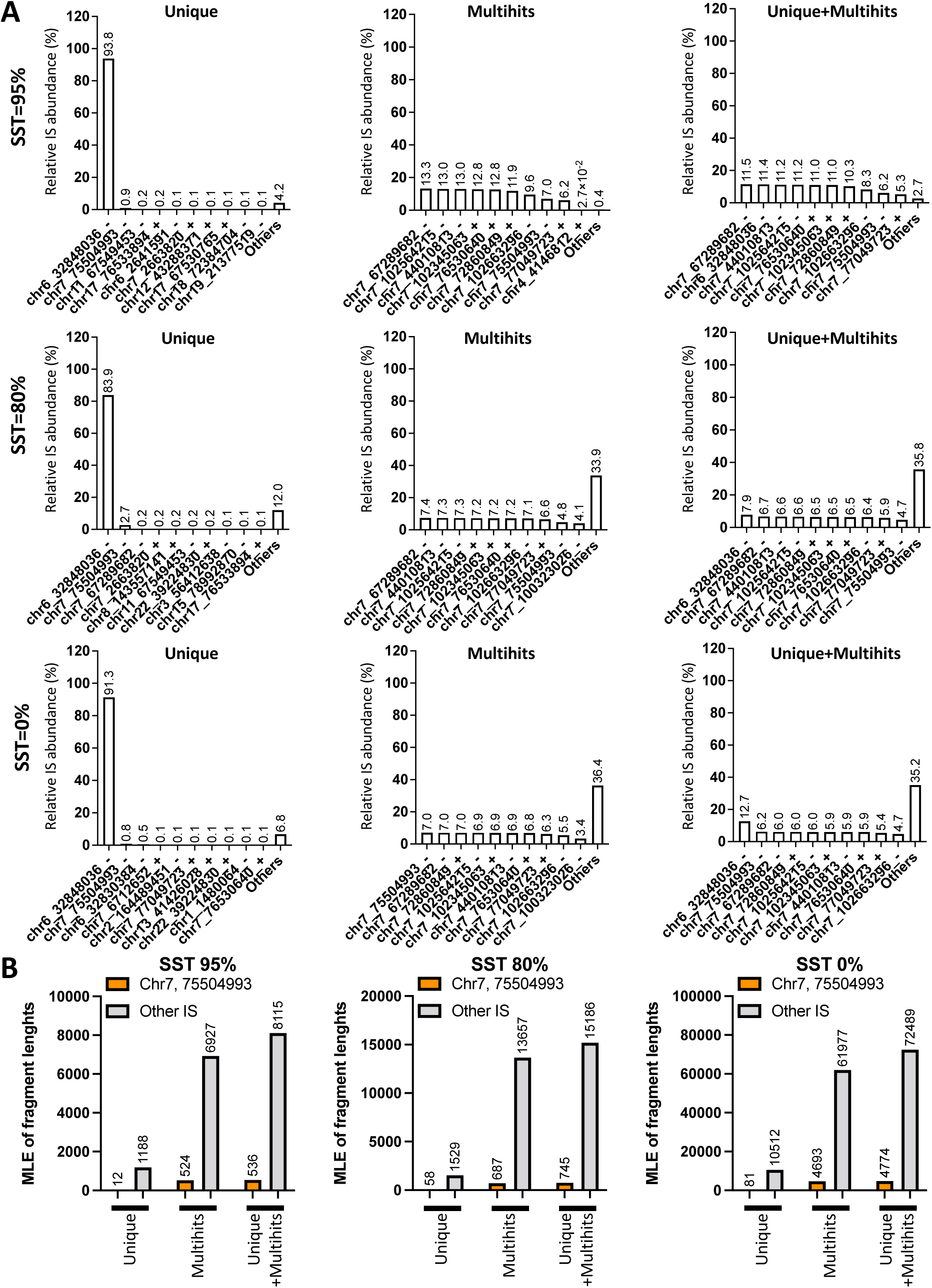
A) Relative abundance of MLE of fragment lengths of IS collected from CLB7 using INSPIIRED unique, multihits and a combination thereof at different sequence similarity thresholds (SST). B) Absolute abundance of the 2^nd^ top IS detected in CLB7 using different SST and different INSPIIRED functions

## Supplementary Tables

**Supplementary Table 1:**
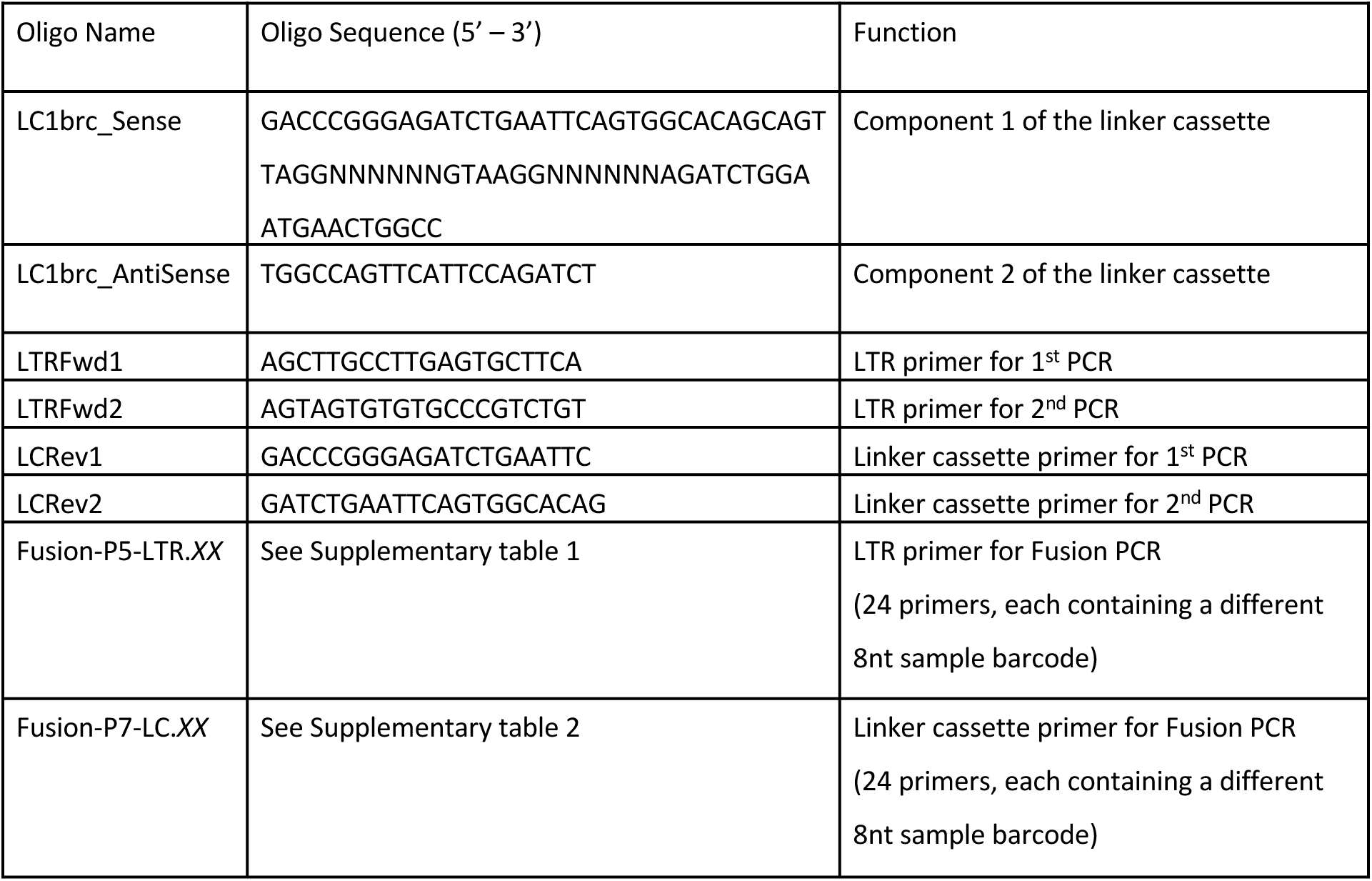
Oligos used for insertion sites retrieval

**Supplementary Table 2:**
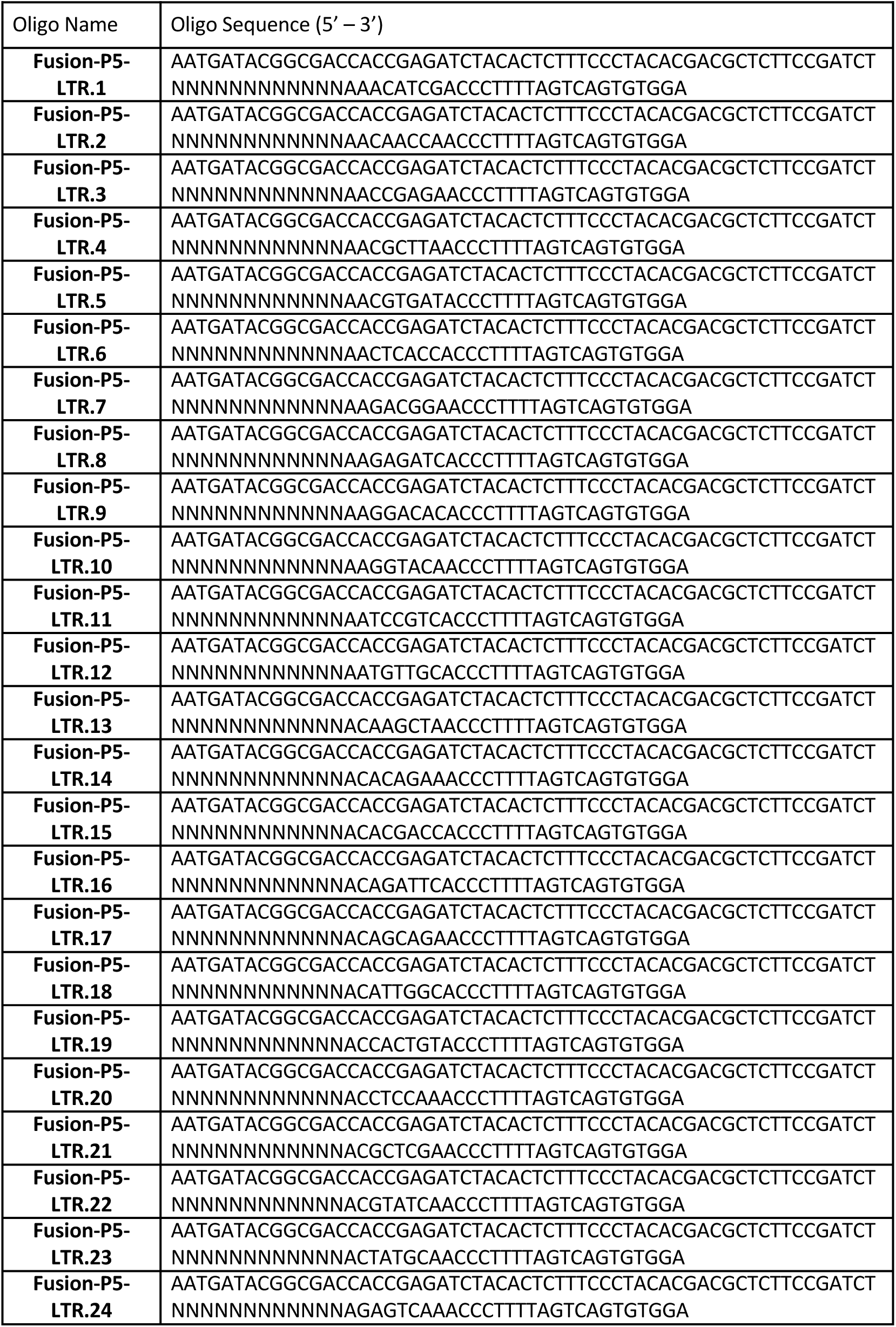
LTR fusion primers used for Illumina sequencing library preparation

**Supplementary Table 3:**
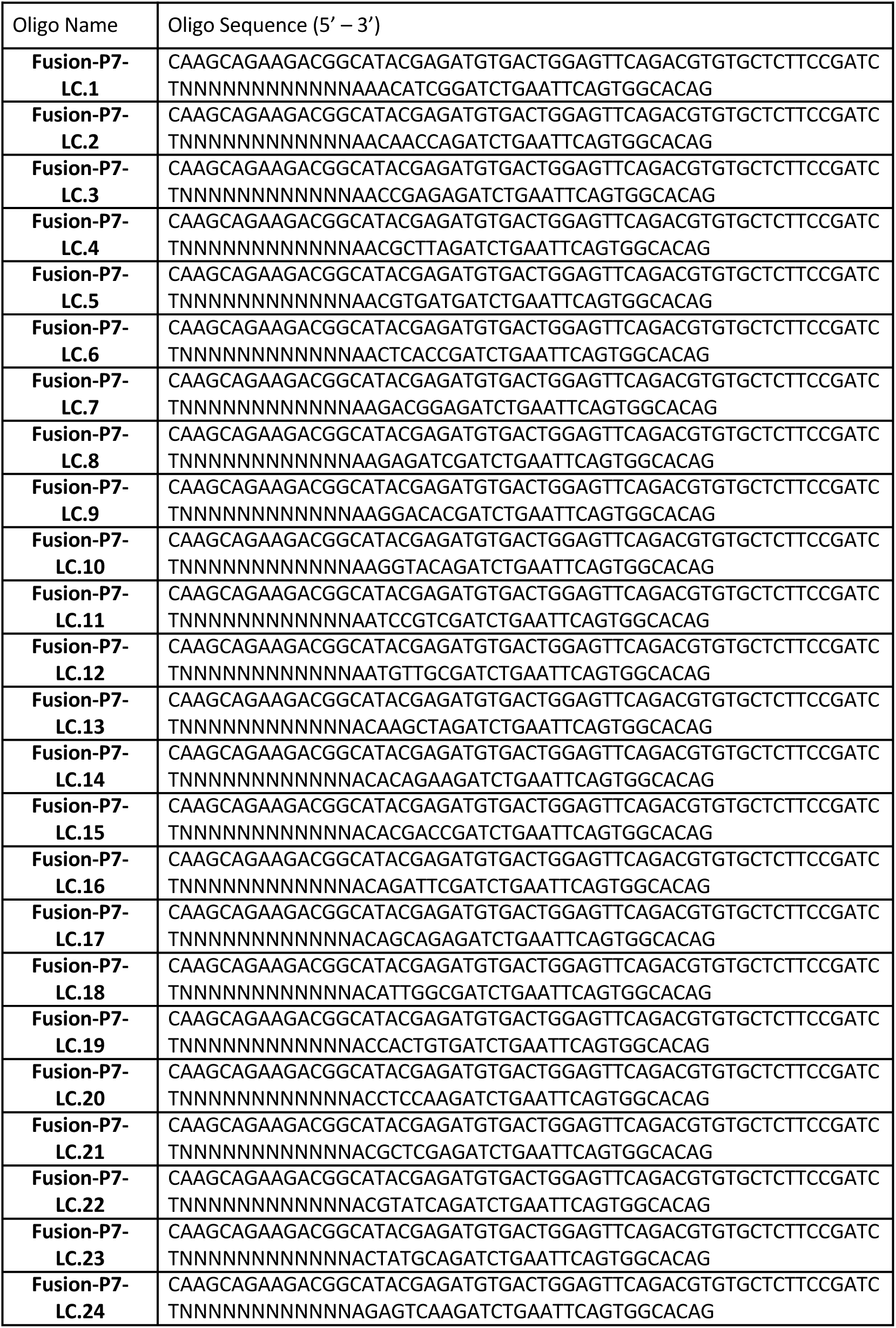
LC fusion primers used for Illumina sequencing library preparation

**Supplementary Table 4:**
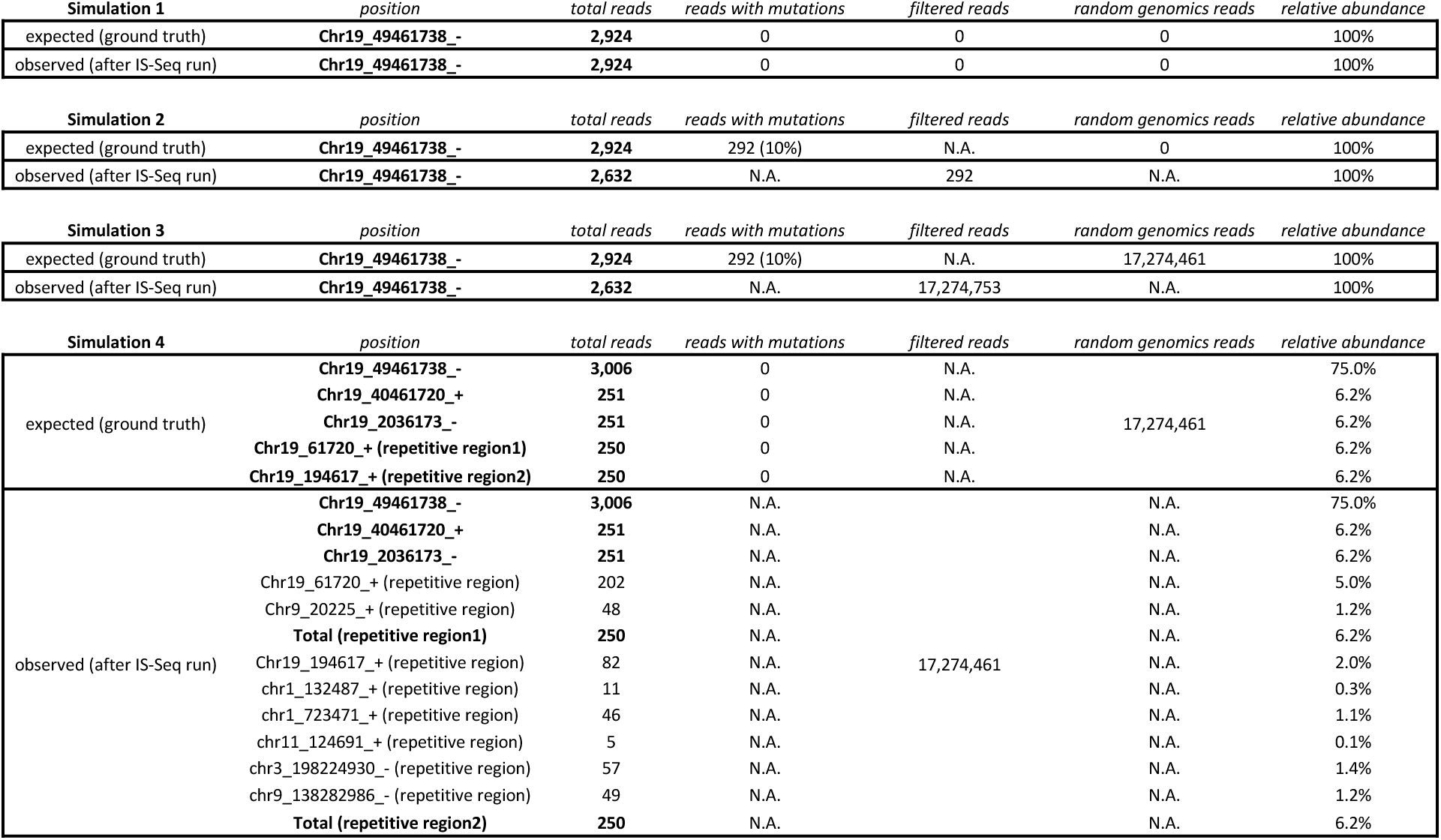
Comparison of expected and observed result from IS-Seq run on 4 *in silico* simulations as described in the manuscript text (N.A= not applicable)

**Supplementary Table 5:** Correlation coefficient (CC) between IS relative abundance calculated with INSPIIRED vs IS-Seq on single cell clones with known copy number.

| Clone ID | CL6 | CLH4 | MOI30<br>CLB6 | MOI30<br>CLB7 | MOI30<br>CLC9 | MOI30<br>CLE1 | MOI50<br>CLB9 | MOI50<br>CLE6 | MOI50<br>CLH6 |
| --- | --- | --- | --- | --- | --- | --- | --- | --- | --- |
| CC | 0.18 | 0.99 | 1.00 | 0.97 | 0.99 | 0.99 | 1.00 | 1.00 | 0.99 |

**Supplementary Table 6:** Mean square error (MSE) calculated between the shared IS resulting from each IS-Seq pipeline iteration (n=2323) and the expected relative abundance of the most abundant IS detected for each clone. The average MSE for all the results of each pipeline iteration is reported in the last row of the table (MLE = Maximum Likelihood Estimation, UMI = Unique Molecular Identifiers)

| Sample Name | VCN | Fragment MLE | Is-Seq Reads | Is-Seq UMI |
| --- | --- | --- | --- | --- |
| CL6 | 1 | 16.48 | 27.99 | 29.98 |
| CLH4 | 5 | 2.90 | 3.09 | 3.12 |
| MOI30CLB6 | 3 | 8.30 | 19.96 | 20.25 |
| MOI30CLB7 | 2 | 722.7 | 94.99 | 92.92 |
| MOI30CLC9 | 2 | 60.7 | 20.8 | 22.2 |
| MOI30CLE1 | 2 | 12.24 | 0.26 | 0.19 |
| MOI50CLB9 | 2 | 5.89 | 0.95 | 0.84 |
| MOI50CLE6 | 2 | 28.34 | 100.73 | 98.88 |
| MOI50CLH6 | 2 | 390.82 | 179.65 | 183.06 |
| Average MSE |  | 138.7078 | 49.82444 | 50.16 |

**Supplementary Table 7:** Mean square error (MSE) calculated between the shared IS resulting from each IS-Seq pipeline iteration and the expected relative abundance of the serial dilutions of CL6 into a bulk HL60 transduced polyclonal population. The average MSE for all the results of each pipeline iteration is reported in the last row of the table (Frag_MLE = Maximum Likelihood Estimation of Fragment Lengths, UMI = Unique Molecular Identifiers)

| IS-Seq | 250<br>(n=1792) |  |  | 500<br>(n=3461) |  |  | 250rep<br>(n=2428) |  |  |
| --- | --- | --- | --- | --- | --- | --- | --- | --- | --- |
| Expected (%) | Frag_MLE | Reads | UMI | Frag_MLE | Reads | UMI | Frag_MLE | Reads | UMI |
| 0 | 0.102407 | 0.0063932 | 0.0065910 | 0 | 0 | 0 | 0.3516624 | 0.0333759 | 0.0347282 |
| 1 | 0.455373 | 0.150538 | 0.143488 | 0.544959 | 0.424843 | 0.433437 | 1.8390805 | 0.7511598 | 0.7521036 |
| 10 | 9.93506 | 5.26387 | 5.16747 | 8.11209 | 7.27422 | 7.13242 | 12.513107 | 7.6450619 | 7.4880557 |
| 50 | 39.6396 | 42.5307 | 42.2344 | 32.3625 | 39.9005 | 39.2183 | 45.820895 | 53.045801 | 52.380017 |
| 75 | 55.8471 | 64.9646 | 64.7314 | 46.0211 | 64.9433 | 64.7753 | 59.679144 | 74.003585 | 73.928068 |
| 100 | 97.0854 | 99.846 | 99.8405 | 99.0991 | 99.9507 | 99.9485 | 88.383045 | 99.423455 | 99.364390 |
| Average MSE | 80.496 | 29.946 | 31.644 | 192.574 | 35.15 | 38.223 | 65.715 | 2.702 | 2.265 |

**Supplementary Table 8:** Number of R2 reads, total UMIs and Unique UMIs derived from the 3 datasets object of this study

| **Data Set** | **Number of R2 reads** | **Number of UMIs extracted from the R2 reads**<br>**(and relative Entropy)** | **Number of unique UMIs extracted from the R2 reads**<br>**(and relative Entropy)** |
| --- | --- | --- | --- |
| Single cell clones Dataset | 17,274,461 | 12,975,641 (22.263)<br>(38% UMI shared by at least 2 reads) | 7,008,854 (22.741) |
| Dilution Dataset 1 (250+500) | 9,102,153 | 5,069,672 (21.888)<br>(17% UMI shared by at least 2 reads) | 4,180,988 (21.995) |
| Dilution Dataset 2 (250rep) | 18,646,287 | 14,422,922 (21.788)<br>(38% UMI shared by at least 2 reads) | 7,107,826 (22.761) |

**Supplementary Table 9:**
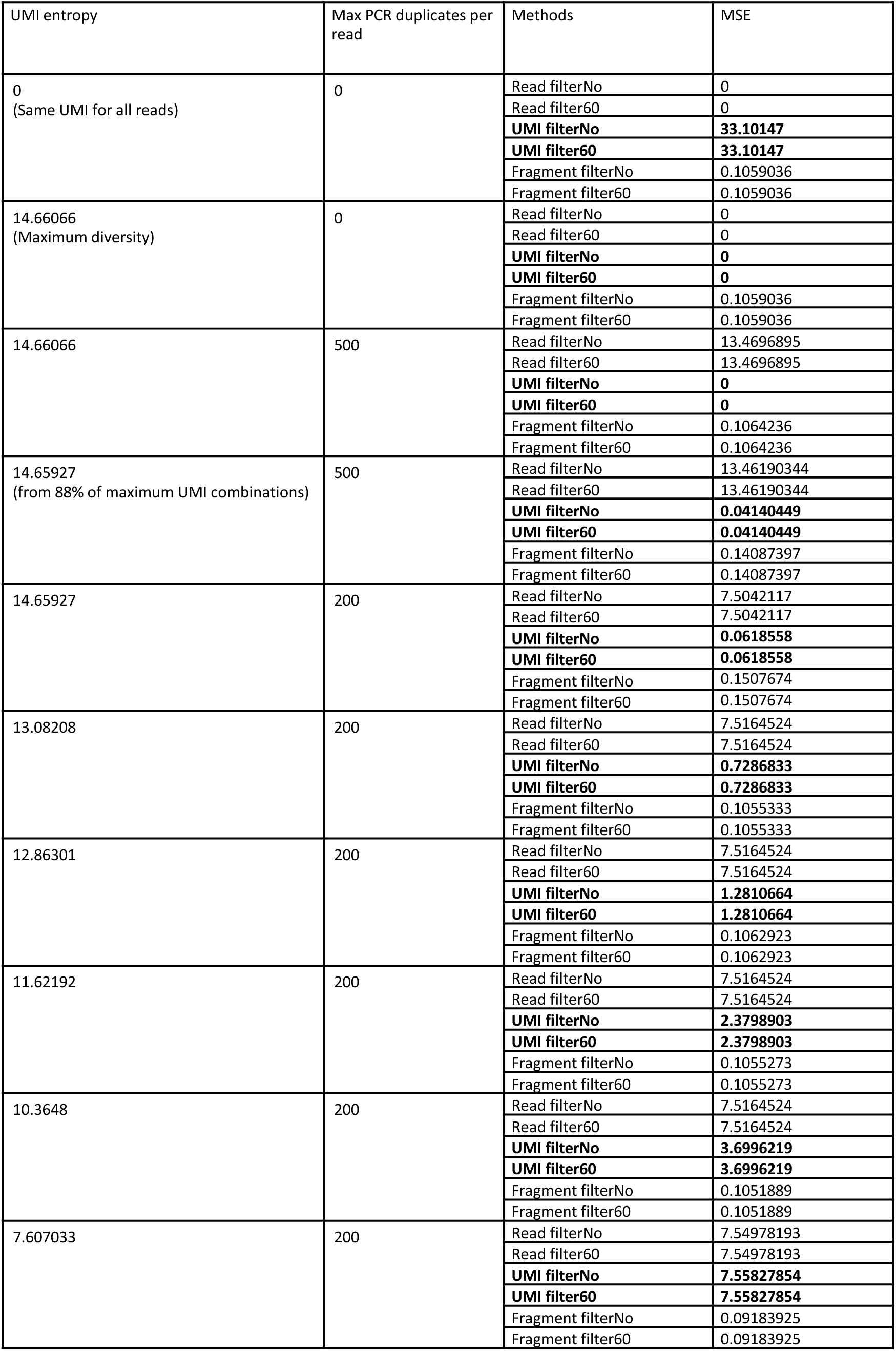
Mean square error (MSE) calculated on a 100 IS simulated dataset with serial IS dilutions using different levels of PCR duplicates and UMI diversity (note: in the simulated dataset FilterNo and Filter60 yield identical results)

